# Tcf21^+^ mesenchymal cells contribute to testis somatic cell development, homeostasis, and regeneration

**DOI:** 10.1101/2020.05.02.074518

**Authors:** Yu-chi Shen, Hailey Larose, Adrienne Niederriter Shami, Lindsay Moritz, Gabriel L. Manske, Qianyi Ma, Xianing Zheng, Meena Sukhwani, Michael Czerwinski, Caleb Sultan, Jourdan Clements, Haolin Chen, Jason R. Spence, Kyle E. Orwig, Michelle Tallquist, Jun Z. Li, Saher Sue Hammoud

## Abstract

Testicular development and function relies on interactions between somatic cells and the germline, but similar to other organs, regenerative capacity decline in aging and disease. Whether the adult testis maintains a reserve progenitor population with repair or regenerative capacity remains uncertain. Here, we characterized a recently identified mouse testis interstitial population expressing the transcription factor Tcf21. We found that Tcf21^+^ cells are bipotential somatic progenitors present in fetal testis and ovary, maintain adult testis homeostasis during aging, and act as reserve somatic progenitors following injury. *In vitro*, Tcf21^+^ cells are multipotent mesenchymal progenitors which form multiple somatic lineages including Leydig and myoid cells. Additionally, Tcf21^+^ cells resemble resident fibroblast populations reported in other organs having roles in tissue homeostasis, fibrosis, and regeneration. Our findings reveal that the testis, like other organs, maintains multipotent mesenchymal progenitors that can be leveraged in development of future therapies for hypoandrogenism and/or infertility.

**Highlights:**

- Multipotent Tcf21^+^ MPs can differentiate into somatic testis cell types
- Tcf21^+^ cells contribute to testis and ovary somatic cells during gonadal development
- Tcf21^+^ cells replenish somatic cells of the aging testis and in response to tissue injury
- Testis Tcf21 cells resemble resident fibroblast populations in multiple organs

## Introduction

Sexual reproduction relies on the generation of distinct sexes to increase biological diversity. The core of this strategy rests with a bipotential gonadal primordium that supports development of sex-specific reproductive organs with functionally distinct gonadal cell types. The gonadal primordium is comprised of primordial germ cells and mesenchymal cells that originate from two sources: the coelomic epithelium (Hatano et al., 1996; Ikeda et al., 1994; Karl and Capel, 1998; Lin et al., 2017; Morohashi, 1997) and the mesonephros (Martineau et al., 1997). In early development, coelomic epithelial cells give rise to multiple cell lineages, including interstitial cells and Sertoli cells (Karl and Capel, 1998), but later become restricted to the interstitial compartment of the testis. In contrast, mesonephric-derived cells migrating into the gonad only contribute to fetal/adult Leydig and interstitial cells, but not Sertoli cells (Buehr et al., 1993; Tilmann and Capel, 1999). Hence, the gonadal mesenchyme is heterogeneous on the molecular and cellular scales, and certain somatic lineages (e.g. Leydig and myoid) have multiple cells of origin (DeFalco et al., 2011), pointing to a complex developmental programming of reproductive organs (reviewed in (Brennan and Capel, 2004; Cederroth et al., 2007; Rotgers et al., 2018).

Establishment of a functional somatic microenvironment is essential for continuous sperm production. In addition to cell types with defined functions, such as testosterone-producing Leydig, contractile myoid, vessel-forming endothelial, immune-response macrophage, and FSH-sensing Sertoli cells, the testis also harbors poorly defined interstitial cell populations (Combes et al., 2009). Emerging studies have highlighted the important roles somatic cells play in germ cell homeostasis and regeneration (Bhang et al., 2018; Green et al., 2018; Kitadate et al., 2019) and how their disruption can dramatically alter testes function. For instance, genetic ablation of macrophages in the adult testis leads to disruption of spermatogonial proliferation and differentiation (DeFalco et al., 2015). Others have shown that testicular endothelial cells produce glial cell line-derived neurotrophic factor (GDNF) as well as other factors that support human and mouse spermatogonial stem cells (SSCs) in long-term culture, and restore spermatogenesis in mice after chemotherapy-induced depletion of SSCs (Bhang et al., 2018). More recently, the Yoshida lab has shown that CD34^+^ interstitial lymphatic endothelial cells secrete FGF, and regulate its niche availability in order to control spermatogonial cell homeostasis and regeneration (Kitadate et al., 2019). Taken together, these studies underscore the importance of defining the interstitial cell composition and function.

Adult interstitial cells are considered postmitotic (not actively cycling) and terminally differentiated. However, studies in rats have demonstrated that adult Leydig cells can regenerate after ethane dimethane sulfonate (EDS)-induced cell death (reviewed in (Smith et al., 2015). Multiple interstitial cell (CD90/PDGFRA/CoupTFII/Nes) populations can re-enter the cell cycle upon EDS-induced Leydig cell death. This suggests that the interstitial compartment contains a reserve Leydig or a general somatic cell progenitor population that can be activated in response to damage (Chen et al., 1996; Chen et al., 2010; Davidoff et al., 2004; Ge et al., 2006; Jiang et al., 2014; Kilcoyne et al., 2014; Kumar and DeFalco, 2018; Landreh et al., 2013; Qin et al., 2008). However, whether the reported molecular markers are expressed universally by a single population of Leydig stem cells or by distinct cell populations that all have the ability to regenerate remains difficult to tease apart. This is due to the difficulty of performing single-cell transplantation and reconstitution experiments in the testis and the lack of multiplexed molecular barcoding tools for lineage tracing (Kumar and DeFalco, 2018; Rotgers et al., 2018; Shima et al., 2018). It is also unclear if a common somatic progenitor could give rise to additional cell types that persists in the adult, and what the role of such stem/progenitors would be in the normal adult testis.

We previously identified a mesenchymal cell population in the adult mouse testis that highly expresses the transcription factor Tcf21 and appears *in vivo* as rare spindle-shaped cells surrounding the seminiferous tubule (Green et al., 2018). The Tcf21 gene has known roles in the development of numerous organs, including the testis (Lu et al., 2000; Lu et al., 2002; Quaggin et al., 1999). Loss of Tcf21 promotes feminization of external genitalia in karyotypically male mice (Cui et al., 2004), while overexpression of Tcf21 in primary embryonic ovary cells leads to *in vitro* sex-reversal via aberrant anti-Mullerian hormone expression (Bhandari et al., 2011). These results provide evidence for a role of Tcf21 in male sex determination and testis somatic cell differentiation. Recent reports of single-cell sequencing during sex fate determination and cell linage specification also identified Tcf21 expression among subsets of gonadal somatic cells in both male and female, although these experiments were limited to NR5A1-egfp cells (Stevant et al., 2019; Stevant et al., 2018). The similarity between the adult Tcf21^+^population to fetal somatic progenitors (Stevant et al., 2019; Stevant et al., 2018) suggests that the Tcf21^+^ population that persist in adulthood may retain fetal developmental functional properties.

In this study, we asked what role Tcf21^+^ cells play in the developing and adult mouse testis. To answer this question, we utilized genetic lineage tracing and a battery of targeted ablation methods to characterize the differentiation potential of Tcf21^+^ cells *in vivo* during fetal gonadal development, adult homeostasis, aging, and injury. We discovered that early Tcf21^+^ progenitors in the fetal gonad contribute to all known somatic cell populations in the adult testis and ovary, with their contribution becoming progressively restricted throughout sexual differentiation and development. We also find that adult Tcf21^+^ cells, persisting from the fetal period, replenish somatic populations in response to injury as well as in normal aging. In directed differentiation paradigms, we find that flow-sorted Tcf21^+^ cells possess mesenchymal stem cell (MSC) like properties and can be directed to differentiate to either myoid or Leydig cell lineages, thus acting as true multipotent progenitors. Furthermore, testis Tcf21^+^ cells resemble resident fibroblast populations in multiple organs. Our work demonstrates the first evidence for a multipotent reserve somatic cell population in the adult testis, representing a potential targetable cell population for development of treatments for gonad-related defects and disease.

## Results

### The Tcf21^+^ population is a mesenchymal cell population that is transcriptionally similar to myoid and Leydig cells

We recently employed single-cell RNAseq (scRNAseq) to generate a molecular atlas of the mouse testis (Green et al., 2018, Sohni et al., 2019)(Shami et al., 2020). Our analysis of 35,000 cells identified not only all somatic cell types known to be present in the adult mouse testis but also an unexpected Tcf21^+^ population (**Figure 1A**) (Green et al., 2018). To better understand its potential function, we first examined if the Tcf21^+^ population has molecular similarity to other known somatic cells in the testis. By using a pairwise dissimilarity matrix for somatic cell type centroids, we find that the Tcf21^+^ population was distinct from macrophage and Sertoli lineages but transcriptionally similar to myoid, Leydig, and endothelial cells (**Figure 1B**). In contrast with these cell types, Tcf21^+^ cells did not express any of their terminally differentiated markers (**Figure 1C**). Rather, this population was uniquely demarcated by the expression of Tcf21 (also known as Pod1), Pdgfra, and mesenchymal progenitor cell (MP) markers including Sca1 (**Figure 1C, S1A; Supplemental Table 1**), Arx, and Vim (Green et al., 2018). To define molecular properties of Tcf21^+^ cells more broadly we identified differentially expressed genes between the Tcf21^+^ population and its most similar cell types (using >2-fold change and FDR <5%). Gene ontology analysis confirmed that the Tcf21^+^ population is of mesenchymal origin, and likely involved in extracellular matrix (ECM) biology, tissue injury and repair processes (**Figure 1D; Table S1**). Although the ECM was once considered a passive support scaffold, a wealth of data has now revealed its active roles in many aspects of biology, from tissue maintenance, regeneration, cell differentiation, to fibrosis and cancer (Bonnans et al., 2014; Hynes, 2009).

**Figure 1:**
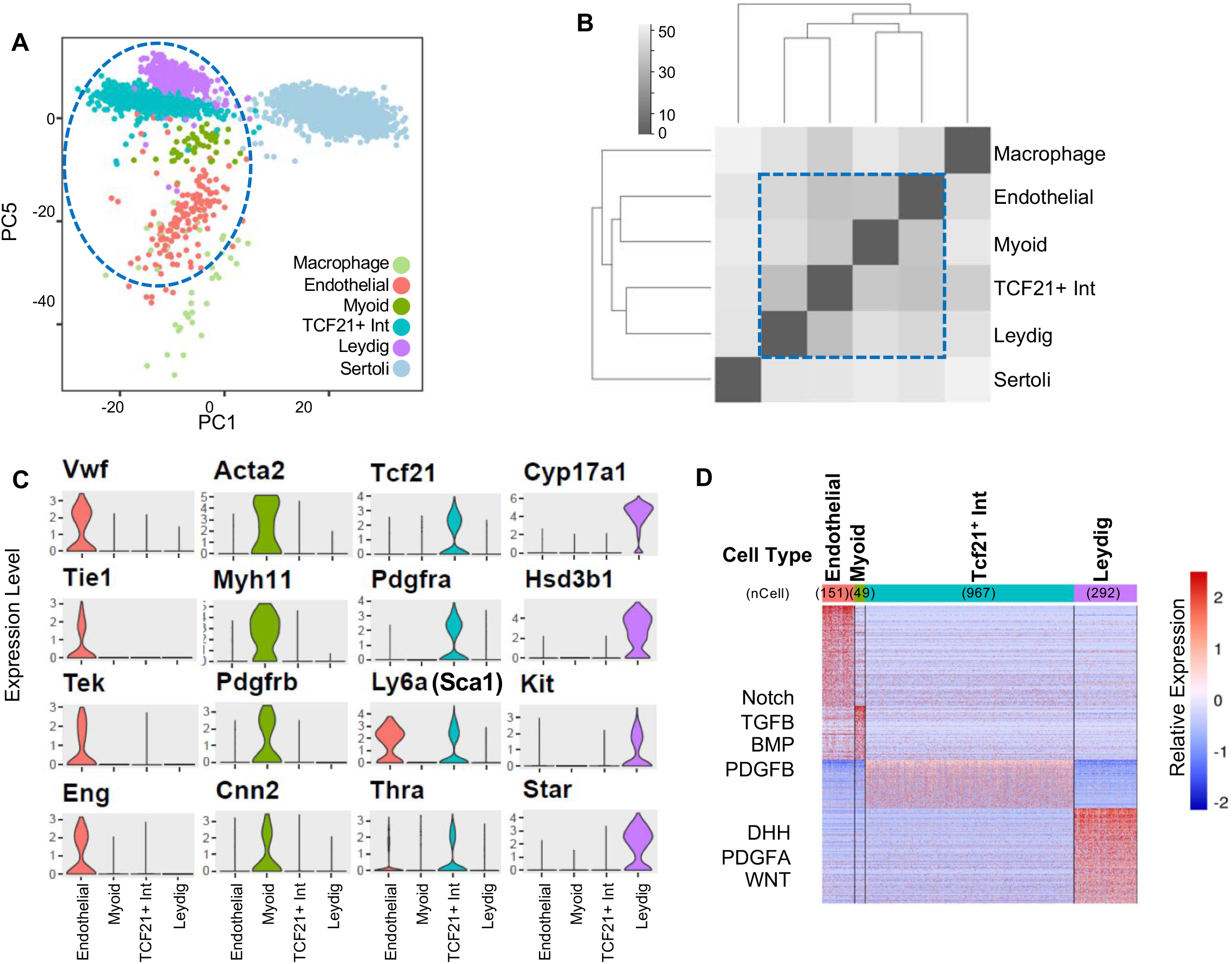
Identification and characterization of a Tcf21-expressing interstitial somatic cell in the adult testis. **(A)** Visualization of 6 somatic cell types in PC space. Data reprocessed from Green et al 2018 – note somatic cell populations were enriched using a combination of cell surface markers or transgenic lines, therefore cell frequencies in PCA plot are not representative of *in vivo*. **(B)** Heatmap of dissimilarity matrix of somatic cell types illustrates high similarity for Tcf21^+^ interstitial population with 3 somatic cell types – myoid, Leydig, and endothelial cells. **(C)** Violin plots of representative markers for endothelial, myoid, Tcf21^+^ interstitial population, and Leydig cells. **(D)** Heatmap of differentially expressed markers for each of the 4 closely related somatic cell types – endothelial, myoid, Tcf21^+^ interstitial population, and Leydig cells.

Given the identification of multiple stromal progenitor markers in the Tcf21^+^ population, we next sought to validate expression of these markers *in vivo*. We took advantage of previously generated *Tcf21^mCrem^* mice, in which a tamoxifen inducible Cre recombinase inserted at the Tcf21 locus enables lineage tracing in the gonads as well in the heart, kidney and cranial muscle (Acharya et al., 2011). Specifically, testes were collected from *Tcf21^mCrem^:R26R^tdTom^* mice after 3 doses of tamoxifen (tdTom^+^ cells are labeled as Tcf21^lin^), dissociated, stained for a comprehensive panel of mouse MSC markers, and analyzed using flow cytometry. Because the Tcf21^lin^ cells were rare in testis histology cross sections (Du et al., 2015; Green et al., 2018), we enriched for Tcf21^lin^ cells using the cell surface marker Sca1 (Ly6a), while excluding mature Leydig (cKit^+^) or immune (CD45^+^) cells (**Figure S1A**). It is important to note that enriching Sca1^+^ cells for scRNA-seq analysis was the approach that originally enabled our discovery of Tcf21^+^ cells. As expected, the Sca1^+^ population has a broader expression pattern in the testis than the specific Tcf21^lin^ (determined by percentage of tdTom^+^ cells), approximately 3-5% vs. 0.72%, respectively (data not shown). Within the Sca1^+^ population, we identified two Tcf21^lin^ subpopulations (blue and purple in **Figure S1B-C**) based on co-expression of mesenchymal stem cell markers: a larger (13.8%, blue) Tcf21^+^ population that strongly expressed CD29 (Itgb1), CD73, and CD34, and a smaller discrete (0.77%, purple) Tcf21^+^ population that expressed all MSC markers. These data suggest that true mesenchymal progenitor cells reside in the latter Tcf21-expressing cell population.

Tcf21^lin^ cells reside near tubules and as determined by our scRNA-seq dataset express several previously described interstitial markers including Pdgfrα, CoupTFII, Cd34, and Fgf5 (Green et al., 2018), we asked whether Tcf21^lin^ could be further defined by co-expression Tcf21^lin^ with several previously reported markers. To answer this question, we performed immunofluorescence analysis of testes from *Tcf21^mCrem^:R26R^tdTom^* or *Tcf21^mCrem^:R26R^tdTom^:PDGFRA^GFP^* male mice co-stained with CoupTFII, Cd34, and FGF5 antibodies (**Figure S1D-G**). Importantly, although our scRNA-seq markers for the Tcf21^+^ cell population included many of the previously reported markers, we did not detect a preferred segregation of Tcf21^lin^ cells within the Pdgfra, CoupTFII, CD34 or Ffg5 subpopulation (**Figure S1D-G**), suggesting that these populations are heterogeneous on both the cellular and molecular level.

### The Tcf21^lin^ /Sca1^+^ population can be differentiated into Leydig and myoid cell fates

Mesenchymal stem cells are typically characterized by their capacity for forming adherent fibroblast-like colonies on plastic and for undergoing adipogenesis, osteogenesis, and chondrogenesis *in vitro* (Reviewed in (Uccelli et al., 2008)). Such populations have been isolated from multiple human and mice organs, including juvenile or adult testes (Ahmed et al., 2017; Eliveld et al., 2019; Gonzalez et al., 2009; Jiang et al., 2014). To determine if and whether the adult Sca1^+^ and Tcf21^lin^ cells have mesenchymal progenitor like properties in vitro, we sorted both Sca1^-^/cKit^+^ interstitial cells (control), Sca1^+^/cKit^-^, or Sca1^+^/Tcf21^lin^ or Tcf21^lin^ cells from *Tcf21^mCrem^:R26R^tdTom^* animals. We found that Sca1^+^ cells as well as Sca1^+^/Tcf21^lin^ cells robustly differentiated into adipocytes, chondrocytes, and osteoblasts (**Figure S2A,B,D**), whereas Sca1^-^/cKit^+^ cells largely fail to thrive and/or differentiate in the trilineage differentiation cocktails (data not shown). Although Sca1^+^, Sca1^+^/Tcf21^lin^, and Tcf21^lin^ populations all formed colonies *in vitro*, clonogenic potential differed across populations. Per 1000 cells plated, cKit^+^ cells (regardless of Sca1 status) formed ~20 colonies, Sca1^+^ cells 30 colonies, and Tcf21^lin^ or Sca1^+^/Tcf21^lin^ cells formed ~100 colonies (**Figure S2C**). Altogether, these data demonstrate that Sca1^+^ and Sca1^+^/Tcf21^lin^ populations have characteristics of mesenchymal progenitors *in vitro*.

Intrigued by the transcriptional relatedness of Tcf21^+^ cells with both Leydig and myoid cells (**Figure 1B**), we next asked whether the Sca1^+^/Tcf21^lin^ population might be a bi-potential progenitor for Leydig and myoid lineages. To answer this question, we performed directed differentiation into Leydig and/or myoid cells. To this end, we sorted Sca1^+^ cells from wild type and *Tcf21^mCrem^:R26R^tdTom^* adult male testes and differentiated these cells using a set of growth factors / cytokines based on the repertoire of receptors expressed in myoid or Leydig cells in our scRNA-seq datasets, and/or based on knowledge previously gained from earlier *in vitro* or genetic experiments (Clark et al., 2000; Li et al., 2016; Tang et al., 2008) (**Figure 2A,B**). After treating Sca1^+^ cells with a myoid differentiation cocktail which includes Smoothened agonist (SAG, an activator of Desert Hedgehog), PdgfAA, PdgfBB, Activin A, Bmp2, Bmp4, and Valproic acid, we observed a morphological conversion of spindle-shaped cells to flattened and striated cells resembling smooth muscle cells (**Figure 2C**). This conversion was confirmed by expression of smooth muscle cell markers such as smooth muscle actin (**Figure 2D, S2D)**.

**Figure 2:**
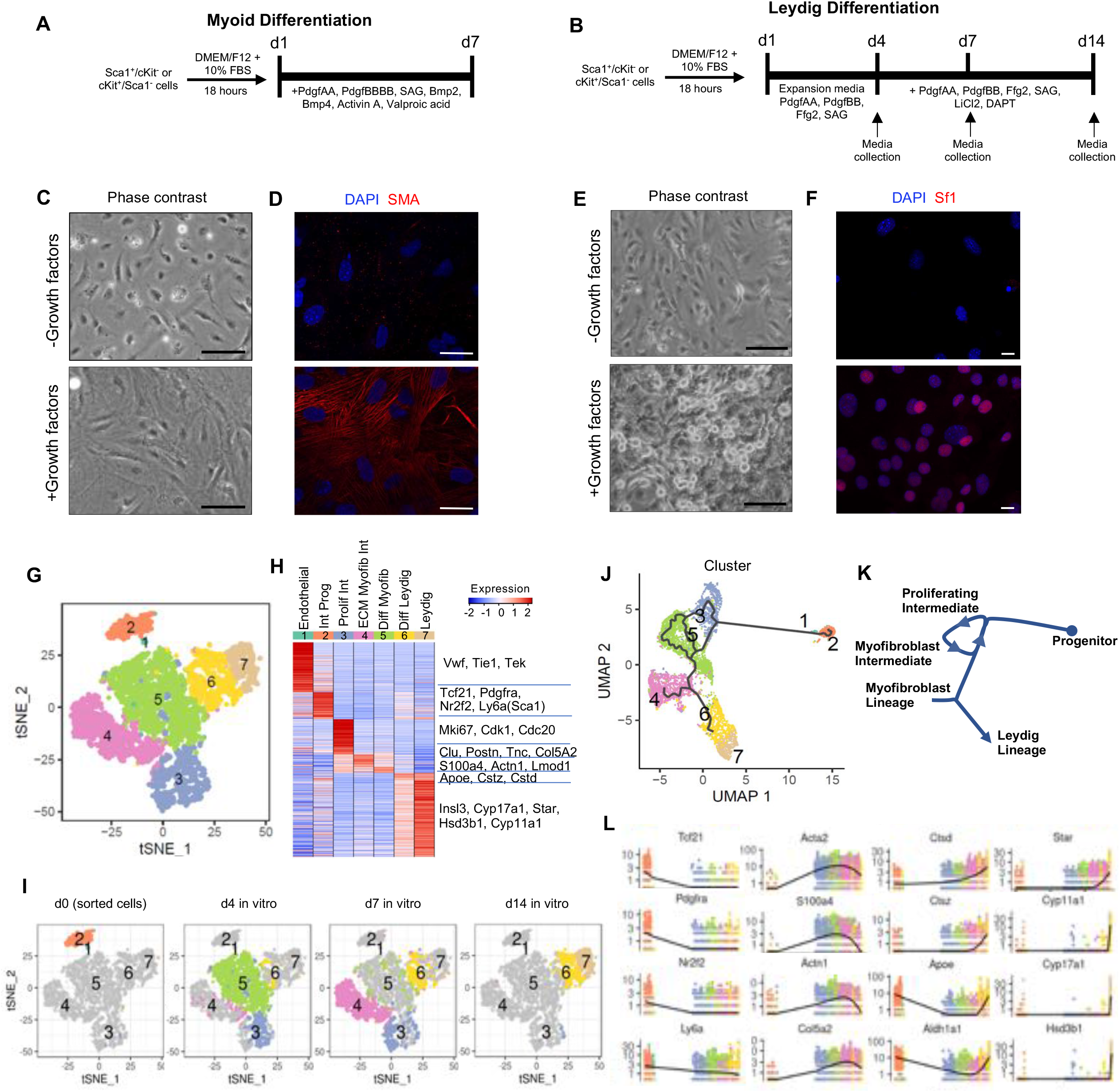
Adult testis Sca1^+^ cells have MSC characteristics and can differentiate to Leydig and smooth muscle cells *in vitro*. **(A-B)** Schematic representation of experimental timelines for myoid **(A)** and Leydig cell **(B)** directed differentiation. **(C-D)** Cell morphology and expression of smooth muscle actin (SMA) in Sca1^+^ cells after 7 days of *in vitro* culture in the presence or absence of a differentiation media via phase contrast **(C)** and immunofluorescence **(D)**. **(E-F)** Cell morphology and expression of steroidogenic factor 1 (SF1) in Sca1^+^ cells after 14 days of *in vitro* culture in the presence or absence of a differentiation media via phase contrast **(E)** and immunofluorescence **(F). (G)** Single cell RNAseq time course analysis of *in vitro* Leydig differentiation (days 0, 4, 7, and 14) identifies seven clusters, as visualized in t-SNE space. **(H)** Heatmap of differentially expressed marker expression levels in the 7 cluster centroids. **(I)** Visualization of the contribution of each individual time point at day 0, 4, 7, or 14 to the 7 clusters in t-SNE space. **(J)** Pseudotemporal ordering of cells from the 4 time points (days 0, 4, 7, and 14) by Monocle3 in UMAP space, colored by 7 clusters. **(K)** Schematic annotation of differentiation trajectory as defined by Monocle. **(L)** Expression profiles of selected markers across the differentiation pseudotime.

Previous experiments in cultured rat seminiferous tubules uncovered that Desert Hedgehog (DHH), FGF, and PDGF signaling are stimulatory to Leydig cell differentiation, while Notch signaling and other factors are inhibitory (Li et al., 2016) (Clark et al., 2000; Tang et al., 2008). Using a similar 14-day differentiation protocol, we found that the majority of adult Leydig cells (Sca1^-^/cKit^+^) were lost in culture by day 3 with only a few cells remaining by day 14 day (data not shown). In contrast, Sca1^+^ and Sca1^+^/Tcf21^lin^ cells expanded and differentiated into Leydig cells after 14 days, and could be maintained for >3 weeks in culture (**Figure 2B,E**). Importantly, the *in vitro*-derived Leydig cells expressed markers of terminally differentiated Leydig cells including steroidogenic factor 1 (SF1) (**Figure 2F, S2E**) and secreted testosterone only in the presence of differentiation cocktail (**Figure S2F-G**). Notably, Leydig cell differentiation and testosterone secretion *in vitro* is independent of LH control, suggesting that the *in vitro*-derived Leydig cells are developmentally locked between fetal Leydig cells (androstenedione secreting but not LH regulated) and adult Leydig cells (LH regulated testosterone secretion (Lei et al., 2001; Ma et al., 2004; Zhang et al., 2001)). Consistent with absence of LH regulation, we detected low levels of luteinizing hormone/choriogonadotropin receptor (LHCGR) expression (~0.48) by day 14 (**Table S2**).

### The Sca1^+^/Tcf21^lin^ population is a multipotent somatic stem cell

Given the Sca1^+^/Tcf21^lin^ population heterogeneity observed by our flow cytometry experiments and co-staining patterns with previously described testis interstitial cell markers, we examined if Sca1^+^/Tcf21^lin^ cells may represent truly multipotent progenitors or are comprised of progenitor subsets that differentiate into different somatic lineages. To test multipotency of Tcf21^lin^ cells directly, we sorted individual Sca1^+^/Tcf21^lin^ into 96-well plates and allowed ~3 weeks of expansion to produce individual clones. Following clonal expansion, clones were directed to differentiate to either myoid or Leydig cell fates. Co-immunostaining with either SMA for myoid cells or SF1 for Leydig cells reveals that individual Sca1^+^/Tcf21^lin^ cells robustly differentiate to adopt either fate with equal frequency (**Figure S2E**), therefore acting as a true multipotent progenitor *in vitro*.

### Single-cell time-course analysis reveals Leydig cell differentiation trajectory

Following the successful directed differentiation of Tcf21^lin^ cells in vitro, we next applied Drop-seq scRNA-seq to molecularly characterize the differentiation process to produce mature Leydig cells. To this end, we collected and analyzed ~6500 cells at four time-points along the *in vitro* differentiation process (d0 Sca1^+^/cKit^-^, d4, d7 and d14 in culture). By clustering the merged dataset and using gene expression and marker gene analysis we identified seven clusters (**Figure 2G**), representing endothelial cells (cluster 1), Tcf21^+^ interstitial progenitors (cluster 2, expressing Tcf21, Pdgfra and CoupTFII), proliferative myofibroblasts (cluster 3), post-mitotic myofibroblasts (clusters 4 and 5), and two subsets of Leydig cells (clusters 6 and 7 enriched for genes involved in steroidogenesis and terminally differentiated Leydig cells) (**Figure 2G-H)**.

When overlaying time points on the different clusters, we find that freshly sorted Sca1^+^/cKit^-^ cells at d0 are present in clusters 1 and 2 (**Figure 2I)**. The small number of cells comprising cluster 1 express von Willebrand factor (VWF) and the endothelium-specific receptor tyrosine kinase Tie-1, suggesting that cluster 1 is a contaminating endothelial cell population from our sorting step (**Figure 2H; Table S2**). Cluster 2 is the interstitial progenitor population which expresses Tcf21, Pdgfra, and CoupTFII (**Figure 2H; Table S2**). Four days after exposure to expansion media, cells dominate in clusters 3 and 5 with a smaller number of cells appearing in clusters 4 and 6 (**Figure 2I)**. Cells in cluster 3 are actively proliferating, as reflected by expression of proliferative marker Mki67, cyclins Cdk1 and Cdc20, and mitotic microtubule associated proteins (**Figure 2H; Table S2**). Cells in cluster 5 are post-mitotic myofibroblasts undergoing dynamic remodeling of cellular organization, and expressing genes involved in actin cytoskeleton dynamics and remodeling (e.g. S100A4, ACTN1, LMOD1, NEXN, ACTA2) (**Figure 2H; Table S2**). Three days (d7) after transition to differentiation media, cells have become directed to an ECM-depositing myofibroblast cell state (expressing: Clu, Postn, Tnc, Col5A2; cluster 4) or a progenitor Leydig cell state (Cluster 6) (**Figure 2H-I; Table S2**). Although cells in cluster 4 do not ultimately contribute to Leydig differentiation, they express fibroblast markers (Col5a2, Postn, Tnc) as well as extracellular-matrix related proteins involved in tissue remodeling (**Figure 2H-I; Table S2**). Interestingly, this population also expresses PDGFA, raising the possibility that Cluster 4 cells serves as an intermediate supportive cell population required to promote continued differentiation of Leydig cells.

By day 14 (10 days of exposure to differentiation media), cells in clusters 6 and 7 have become more differentiated (**Figure 2I**). Cells in cluster 6 appear to prepare for steroidogenesis by increasing expression of lysosome/exosome genes, likely employing autophagy to degrade cellular components into steroid building blocks like cholesterol and fats (**Figure 2H; Table S2**). Previously, autophagy in Leydig cells was shown to be a rate-limiting step for testosterone synthesis (Gao et al., 2018). Apolipoprotein E (ApoE) is also expressed at this time, indicating that LDL uptake, critical for steroidogenesis, is occurring, in line with genetic evidence in ApoE/LDLR knockout mice (Steinfeld et al., 2018). Finally, in cluster 7, steroidogenic enzymes CYP17A1, HSD3B1, StAR, CYP11A1 are expressed, as well as the mature Leydig cell factor INSL3, indicating a cellular state with functional steroidogenesis (**Figure 2H; Table S2**). Our *in vitro* developmental progression based on time point sampling was confirmed by Monocle3 pseudotime analysis (**Figure 2J-L**), where sorted progenitors give rise to a cycle of proliferating and differentiating intermediates. Cells then branch into two differentiation trajectories, one aborting in cluster 4, which is an ECM producing myofibroblast-possibly a support intermediate cell, while the remaining cells proceed through clusters 6 and 7 which lead to differentiated Leydig cells.

Given that Leydig cell differentiation and testosterone secretion *in vitro* was LH independent, we asked whether the *in vitro* derived Leydig cells resemble the fetal or adult Leydig cell states. To explore this, we performed cluster-cluster correlations between the *in vitro* defined molecular states to previously published adult and fetal *in vivo* somatic cell states (Green et al., 2018; Stevant et al., 2018). Notably, the *in vitro* intermediate states (Clusters 2-5) in Leydig cell differentiation correlate with early interstitial progenitors in the fetal gonad, whereas, clusters 6 and 7 have a higher correlation to fetal Leydig cells (**Figure S2G**). When comparing to the adult testis, the *in vitro* intermediate states (Clusters 2-5) correlate with the Tcf21-expressing interstitial population, but clusters 6 and 7 have the highest correlation to adult Leydig cells (**Figure S2H)**. Interestingly, the correlation is much higher for adult Leydig cells than fetal Leydig cells, suggesting that despite the lack of LH regulation in vitro, the in vitro derived Leydig cells are transcriptionally more similar to adult than fetal Leydig cells (**Figure S2H,I**).

### Tcf21^lin^ cells contribute to somatic lineages in the male gonad

Our previous work showed a high transcriptional correlation between the adult Tcf21-expressing population and a fetal somatic interstitial progenitor population (Green et al., 2018), which we also confirmed with a recent dataset from the developing testis (**Figure S2I**). Given these data, we next asked if the Tcf21 lineage can serve as a somatic progenitor *in vivo*. To this end, we performed lineage-tracing from early developmental time points in *Tcf21^mCrem^:R26R^tdTom^* mice and analyzed fully formed testes at E17.5 and adult testes at 10 weeks. Specifically, timed pregnant females were injected with a single dose of tamoxifen at gestational days E9.5, E10.5, E11.5 and E12.5 (**Figure 3A**). We verified that Tcf21^lin^ labeling was absent in embryonic gonads harvested from vehicle-treated timed pregnant females, confirming tight regulation of the tamoxifen inducible Cre (**Figure S3A**). Additionally, lineage traced cells did not overlap with Vasa, a germ cell marker, confirming specificity of the *Tcf21^mCrem^* Cre line (**Figure S3B)**.

**Figure 3:**
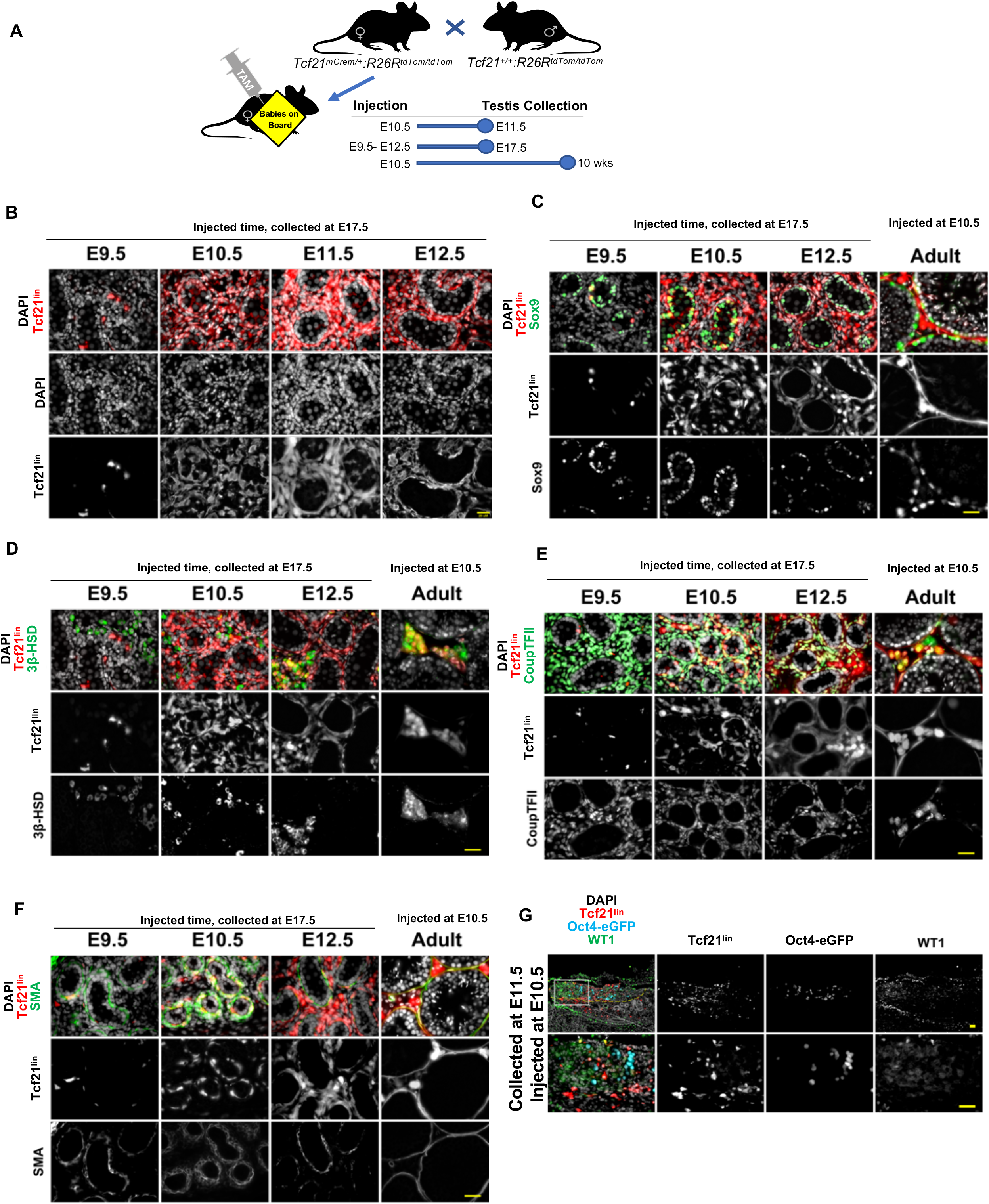
The Tcf21^lin^ population contributes to multiple somatic lineages in the fetal and adult testis. **(A)** Experimental timeline used for Tcf21 lineage tracing analysis. *Tcf21^mCrem^:R26R^tdTom^* timed pregnant females were injected with a single dose of Tamoxifen at E9.5, 10.5, E11.5 or 12.5 and testes were analyzed at E17.5 or 10 weeks. **(B)** The Tcf21^lin^ cells at E10.5 and E11.5 contribute to all major somatic cell populations in the fetal gonad, whereas the E12.5 Tcf21^lin^ cells give rise to testis interstitial cells. **(C-F)** Co-immunostaining of fetal or fostered adult Tcf21^lin^ testis cross-sections with Sertoli cell marker Sox9 **(green; C),** Leydig cell marker 3βHSD **(green; D)**, interstitial cell marker CoupTFII **(Nr2F2; green in E),** and a myoid cell marker alpha smooth muscle actin **(SMA, green in F)**. (**G**) Exploring the origin of Tcf21^lin^ in the fetal testis of *Tcf21^mCrem^; R26R^tdTom^;* Oct4-eGFP embryos. Colocalization of WT1+ **(Green)** cells in the E11.5 gonads withTcf21^lin+^ cells. In all panels the nuclear counterstain is DAPI **(white B-G)**. Scale bars: 20μm.

Immunofluorescence and histological analysis of E17.5 male gonads revealed marked differences in the extent of labeling across the different tamoxifen injection time points (**Figure 3B**). We observed relatively fewer Tcf21^lin^ cells from E9.5 injected animals, yet those cells colocalized with multiple somatic lineages including Sertoli (SOX9), fetal Leydig cells (3BHSD), myoid (SMA) cells, and interstitial cell markers (CoupTFII) (**Figure 3C-F, S3C-F)**, suggesting that early injections are labeling very few early Tcf21 cells that are multipotent. However, the overwhelming majority of Tcf21^lin^ cells contributed to Sertoli cells (**Figure S3C-F)**. In contrast, a greater number of Tcf21^lin^ cells are observed in the E17.5 gonads collected from animals treated with tamoxifen at E10.5, E11.5 and E12.5, possibly due to broader expression of Tcf21 in multiple somatic progenitors (**Figure 3C-F, S3C-F)**. Nevertheless, at these time points, we found that the Tcf21^lin^ cells again contributed to Sertoli (SOX9), fetal Leydig (3BHSD), interstitial (CoupTFII, PDGFRA, and GLI), and myoid (SMA) cells (**Figure 3C-F, S3I-H**). However, the Tcf21^lin^ contribution to the Sertoli lineage became progressively restricted from E10.5 and on, suggesting that developmental competency of the Tcf21 population is restricted or limited by the Sertoli cell specification window.

Early *in vitro* lineage tracing experiments using phospholipid dyes enabled the discovery that cells migrating from the coelomic epithelium can give rise to all somatic cells including Sertoli and interstitial populations in the testis (Karl and Capel, 1998). However, these approaches could not reveal the molecular identity of the cells. Subsequent related work by others reported that a WT1^+^ (Wilms’ tumor 1) somatic progenitor population is essential for gonadogenesis (Armstrong et al., 1993; Kreidberg et al., 1993; Liu et al., 2016) and the WT1 lineage tracing experiments demonstrated that the WT1^+^ population gives rise to Sertoli and interstitial populations including adult Leydig cells (Liu et al., 2016). In our next set of experiments, we set out to determine the origin of Tcf21^+^ cells in the embryonic gonad and potential overlap with WT1^+^ cells. To achieve this goal, we crossed *Tcf21^mCrem^:R26R^tdT^* mice with an *Oct4-eGFP* transgenic mouse line (Szabo et al., 2002) to facilitate visualization of primordial germ cells and identification of the developing bipotential gonad. We then injected timed pregnant *Tcf21^mCrem^:R26R^tdT^;Oct4-eGFP* females with a single dose of tamoxifen at E10.5 and collected gonads at E11.5. In E10.5 injected animals, we found that Tcf21^lin^ cells localize to both the gonad/coelomic epithelium and the mesonephros, making it difficult to determine if Tcf21^lin^ cells truly originate from coelomic or mesonephros, or are present in either location. However, we find that Tcf21^lin^ cells partially overlap with the WT1 population, but many cells are either Tcf21^lin^ or WT1^+^ (**Figure 3G**).

Of the somatic cell populations in the testis, the steroid producing Leydig cells are unique in that they arise in two distinct waves. Fetal Leydig cells (FLCs) develop in the mammalian testis prenatally and are essential for masculinization of fetal tissues, but are then postnatally replaced by adult Leydig cells (ALCs), which are responsible for secondary sex characteristics and the maintenance of spermatogenesis (Roosen-Runge and Anderson, 1959). To ascertain whether fetal-derived Tcf21^+^ lineage persist in the postnatal testis and give rise to adult Leydig cells, the *Tcf21^mCrem^:R26R^tdT^* time pregnant females received a single injection of tamoxifen at E10.5 and the pups were fostered and matured to adulthood (**see Methods; Figure 3A**). In 10-week old male testes, we found that a fraction of adult Sertoli, peritubular myoid, Pericytes or endothelial cells, and interstitial cells are tdTom^+^, suggesting that these somatic populations originated from the fetal Tcf21^lin^ population. However, similar to the embryonic gonad, tdTom is detected in a fraction of the different cell types (**Figure 3C-F)** consistent with the notion of multiple cells of origin or an unlikely possibility of incomplete tamoxifen labeling of embryonic cells. Furthermore, we observe overlap between Tcf21^lin^ and CoupTFII or Tcf21^lin^ and 3BHSD, suggesting that the fetal Tcf21^lin^ population persists in the interstitial space of the postnatal testis and gives rise to adult Leydig cells **(Figure 3D,E).**

Overall, these results suggest that the fetal Tcf21^lin^ *in vivo* is either a multipotent or molecularly heterogenous population of somatic progenitor populations that gives rise to most differentiated somatic populations present in the testis, including fetal Leydig cells, adult Leydig cells, peritubular myoid cells, vascular endothelial cells, and Sertoli cells. While the Tcf21^lin^ population overlaps with multiple previously described interstitial markers (CoupTFII, WT1, GLI, PDGFRA) present during testis development and specification of somatic cells, the Tcf21^+^ population represents a separate, yet parallel early pool of progenitors in gonadal development.

### The Tcf21^lin^ give rise to multiple fetal and adult ovarian somatic cell types

Prior to sex determination in mammals, the gonadal primordium is bipotential, meaning that the gonadal mesenchyme has the ability to give rise to either male or female support cell and steroidogenic cell lineages (Stevant et al., 2019). Once Sry expression is turned on during a critical window of fetal development, testis differentiation is initiated (Koopman et al., 1990). Despite an early commitment to either male or female somatic cell types, genetic studies in mouse have shown that terminally differentiated cell types must be actively maintained throughout life. For example, loss of either *Dmrt1* in Sertoli cells or *Foxl2* in granulosa cells can trigger reciprocal cell fate conversions (from Sertoli to granulosa cell fate or granulosa to Sertoli cell fate, respectively, (Matson et al., 2011; Matson and Zarkower, 2012; Uhlenhaut et al., 2009). Taking into account the known gonadal mesenchyme plasticity and the ability of Tcf21^lin^ cells to give rise to multiple somatic lineages in the testis, we next asked whether Tcf21^lin^ cells are present in the fetal ovary and whether they might give rise to analogous cell types in females. For our female gonad experiments, we injected *Tcf21^mCrem^:R26R^tdT^;Oct4-eGFP* mice with a single dose of tamoxifen at E10.5 and collected female embryonic gonads the next day E11.5. Our analysis shows that Tcf21^lin^ cells are present in the coelomic epithelium, mesonephros and gonads, similarly to what we had observed in male embryonic gonads (**Figure S3K**). We then injected tamoxifen in timed-pregnant females at E10.5, E11.5, or E12.5 *Tcf21^mCrem^:R26R^tdT^* mice and found broad somatic cell labeling in the female gonads at E19.5 (**Figure S3L)**. Co-staining ovarian cross-sections with terminally differentiated markers shows that Tcf21^lin^ cells overlap with granulosa cell marker (FOXL2^+^), interstitial cells (CoupTFII^+^), and smooth muscle cells (SMA^+^), and does not overlap with the germ cell markers Vasa or Oct4 (**Figure S3M-P)**.

In the mouse ovary, androgen producing theca cells are not morphologically distinguishable one week after birth but are easy to identify once they surround the follicles (Edson et al., 2009). However, specification of theca cells has already occurred around the time of birth, and is postnatally derived from Gli1^+^ and/or WT1^+^ populations (Honda et al., 2007; Liu et al., 2015). We asked if the embryonic Tcf21^lin^ cells overlap with WT1^+^ progenitors and if fetal Tcf21^lin^ cells give rise to theca cells in the postnatal ovary. To this end, we examined E11.5 embryonic gonads from timed pregnant *Tcf21^mCrem^:R26R^tdT^;Oct4-eGFP* mice and co-stained gonad sections for WT1 (**Figure S3Q**). WT1 and Tcf21^lin^ had little overlap indicating that WT1 and Tcf21^lin^ mark largely distinct populations not originating from the same progenitor (**Figure S3Q**). However, when we traced the fate of fetal (E10.5) Tcf21^lin^ cells in the adult ovary, we find that the embryonic Tcf21^lin^ population contributes to granulosa cells (FOXL2^+^ and WT1^+^), endothelial cells (Pecam^+^), and theca cells (3BHSD^+^), but not germ cells (**Figure S3Q)**. Based on these data, we conclude that the fetal Tcf21^+^ population is likely also a multipotent somatic progenitor in the female.

### Tcf21^lin^ cells regenerate somatic cell types in the adult mouse testis

The somatic cells of the testis are considered to be postmitotic and do not naturally turnover. However, in our steady state scRNA-seq datasets, we found rare cells that cluster within the Tcf21^+^ interstitial population that express low levels of either steroidogenic acute regulatory protein (StAR) or SMA, suggesting that a very rare population of Tcf21^+^ cells are differentiating into either Leydig or myoid cells, respectively. In our next set of experiments, we asked if the adult Tcf21 lineage is capable of regenerating somatic testis cells. To answer this question, we set up genetic and chemical ablation models to reduce Leydig and myoid cell numbers and assess Tcf21^lin^-mediated regeneration of these populations.

To genetically ablate Leydig cells, we crossed Rosa26^iDTR^ and Cyp17a1-icre mice (Bridges et al., 2008) to generate *Cyp17a1-cre; Rosa26*^*iDTR*/+^ mice (hereafter referred to as *Cyp17a1-Cre; iDTR*). At the age of 8 weeks, male mice received four serial injections of diphtheria toxin (DTX), titrated to achieve maximum cell death while ensuring animal survival (**Figure 4A**). Based on the timeline for Leydig cell regeneration in rats after chemical ablation (reviewed in (Smith et al., 2015), we collected mouse testes at 24 hours post final injection (24hpfi), 3 dpfi, 7 dpfi, 14 dpfi, and 21 dpfi. In the genetic model, Leydig cells undergo apoptosis within 24 hours (**Figure 4B-C**), and concomitantly testosterone levels drop and recover by ~21 days (**Figure 4D-E**). Although the genetic approach works with high consistency, the majority of animals (~98%) do not survive the duration of the experiment.

**Figure 4:**
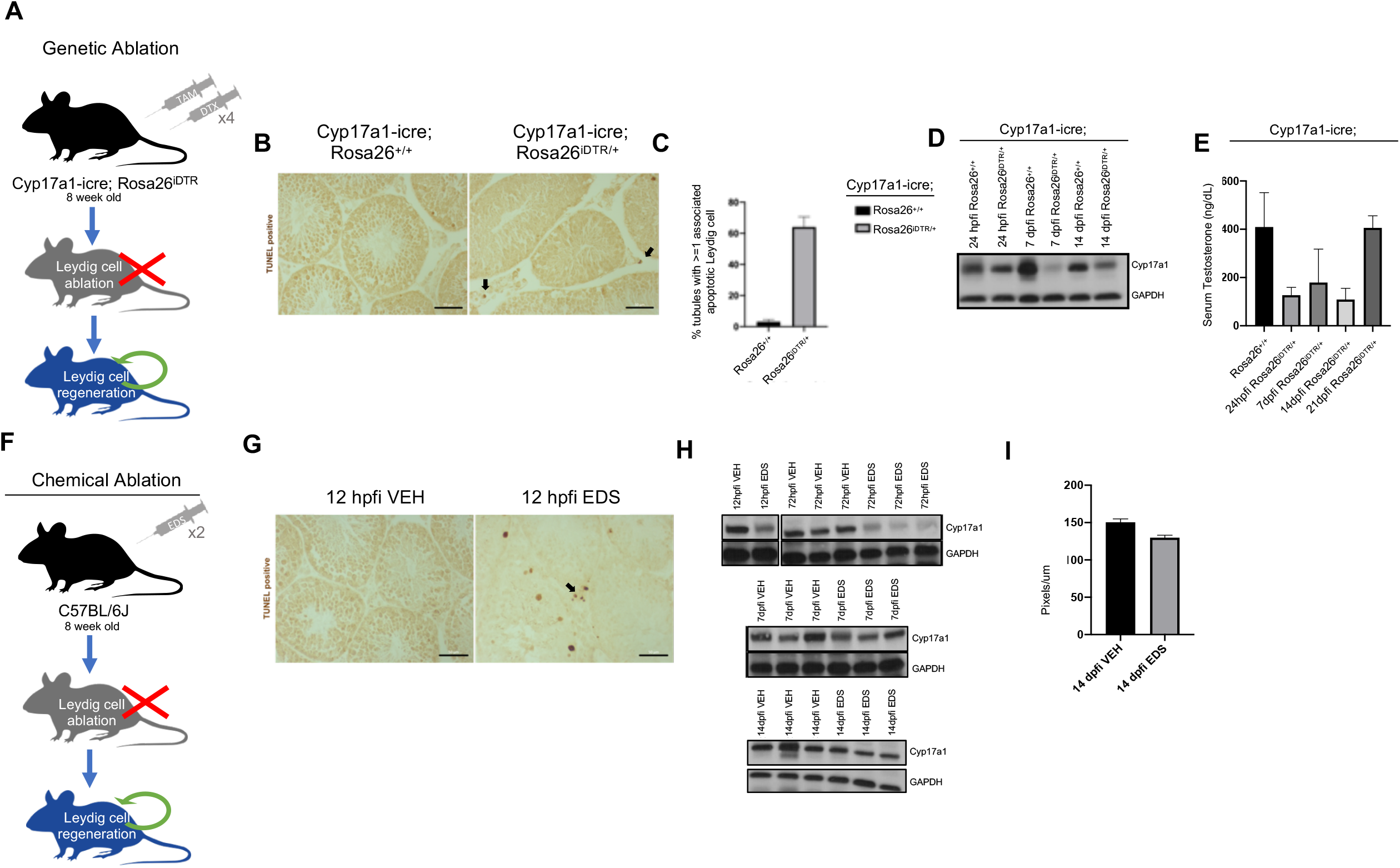
Chemical and genetic approaches ablate Leydig cells. **(A)** Schematic representation of the genetic method used to ablate Leydig cells. **(B-F)** Efficacy of diphtheria toxin (DTX) treatment in *Cyp17a1-icre;Rosa*^*DTR*/+^ or *Cyp17a1-icre;Rosa*^+/+^ (wild type) animals. **(B)** Representative images of apoptotic Leydig cell ablation by TUNEL staining in testes collected 24 hours after final injection (hpfi) of DTX in *Cyp17a1-icre;Rosa*^*DTR*/+^ or *Cyp17a1-icre;Rosa*^+/+^. **(C)** Quantification of tubules containing at least one associated apoptotic Leydig cell in *Cyp17a1-icre;Rosa*^*DTR*/+^ or *Cyp17a1-icre;Rosa*^+/+^ mice 24hpfi of DTX in at least 50 randomly selected tubules per animal (n=2). **(D)** Representative Western Blot illustrating Cyp17a1 protein expression levels at 24hpfi, 7dpfi, and 14dpfi in *Cyp17a1-icre;Rosa*^*DTR*/+^ or *Cyp17a1-icre;Rosa*^+/+^ animals. **(E)** Serum testosterone measurements at 24hpfi, 7dpfi, 14dpfi and 21dpfi in *Cyp17a1-icre;Rosa*^*DTR*/+^ or *Cyp17a1-icre;Rosa*^+/+^ animals (n=2). **(F)** Schematic representation of the chemical method used to ablate Leydig cells. **(G)** Representative images of apoptotic Leydig cells by TUNEL staining in testes collected 12hpfi injection of EDS. **(H)** Western Blot illustrating Cyp17a1 protein expression levels at 12hpfi, 72hpfi, 7dpfi, and 14dpfi in *C57BL/6* animals after EDS or vehicle (VEH) treatment (n=3). **(I)** Average Leydig cell diameter measurements 14dpfi of EDS or vehicle in *C57BL/6* animals, after Leydig cell recovery. At least 80 cells were measured per animal (n=3); Scale bars: 20 μm.

We therefore next turned to chemical ablation with ethane dimethane sulfonate (EDS) (**Figure 4F**), established in rats to selectively ablate Leydig cells within the testes (Jackson, 1970; Kerr et al., 1985; Molenaar et al., 1986; Morris et al., 1986). Although there is strong species specificity and animal-to-animal differences in Leydig cell sensitivity to EDS (Ewing and Keeney, 1993), we found that two 300 mg/kg EDS injections in C57BL/6 mice spaced 48 hours apart resulted in significant reduction in mature Leydig cells as evidenced by cell death and Cyp17a1 protein levels (**Figure 4G**). At 12hpfi, we detected TUNEL positive Leydig cells, but given the spacing of 48 hours between the first and second injection, the majority of apoptotic events preceded the time of analysis and could therefore not be quantified (**Figure 4G**). However, consistent with Leydig cell loss, we find that Cyp17a1 protein expression decreased as early as 12hpfi and had largely recovered by 14dpfi (**Figure 4H**). We confirmed that the recovery of Cyp17a1 protein expression was not simply due to Leydig cell hypertrophy, as the Leydig cell diameter is similar across *Cyp17a1-cre; Rosa26^iDTR/+^* and *Cyp17a1-cre; Rosa26^+/+^* animals at 14dpfi (**Figure 4I**).

To determine if regenerating Leydig cells were derived from the adult testis Tcf21^lin^ we labeled Tcf21^+^ cells in adult *Tcf21^mCrem^:R26R^tdTom^* mice with three 2 mg tamoxifen injections, and then treated the mice with two 300mg/kg EDS injections spaced 48 hours apart (**Figure 5A**). Testes were collected at 24hpfi, 3dpfi and 24dpfi to assay Leydig cell death, Tcf21^lin^ cell proliferation, and Leydig cell regeneration. Treating the *Tcf21^mCrem^:R26R^tdTom^* mice with EDS followed similar dynamics as in C57BL/6 animals. Leydig cell number decreased at 3dpfi and recovered by 24dpfi, as evidenced by comparing Cyp17a1 protein levels in EDS-treated and vehicle-treated animals (**Figure 5B**). At 3dpfi, the Tcf21^lin^ cells surrounding the tubules reentered the cell cycle as detected by co-localization of BrdU and Tcf21^lin^ (**Figure 5C**). Importantly, by 24dpfi in EDS-treated animals a fraction of Leydig cells, marked by SF1, were Tcf21^lin^ positive, whereas this was not the case in either vehicle treated animals or in EDS-treated mice at 24hpfi (**Figure 5D**). As in chemically ablated C57BL/6 mice, we found that the Tcf21^lin^/SF1^+^ cells have a similar diameter to SF1^+^ cells (**Figure 5E**). Therefore, we demonstrate that the adult Tcf21^lin^ population in the testis can serve as a reserve Leydig progenitor in response to Leydig cell loss.

**Figure 5:**
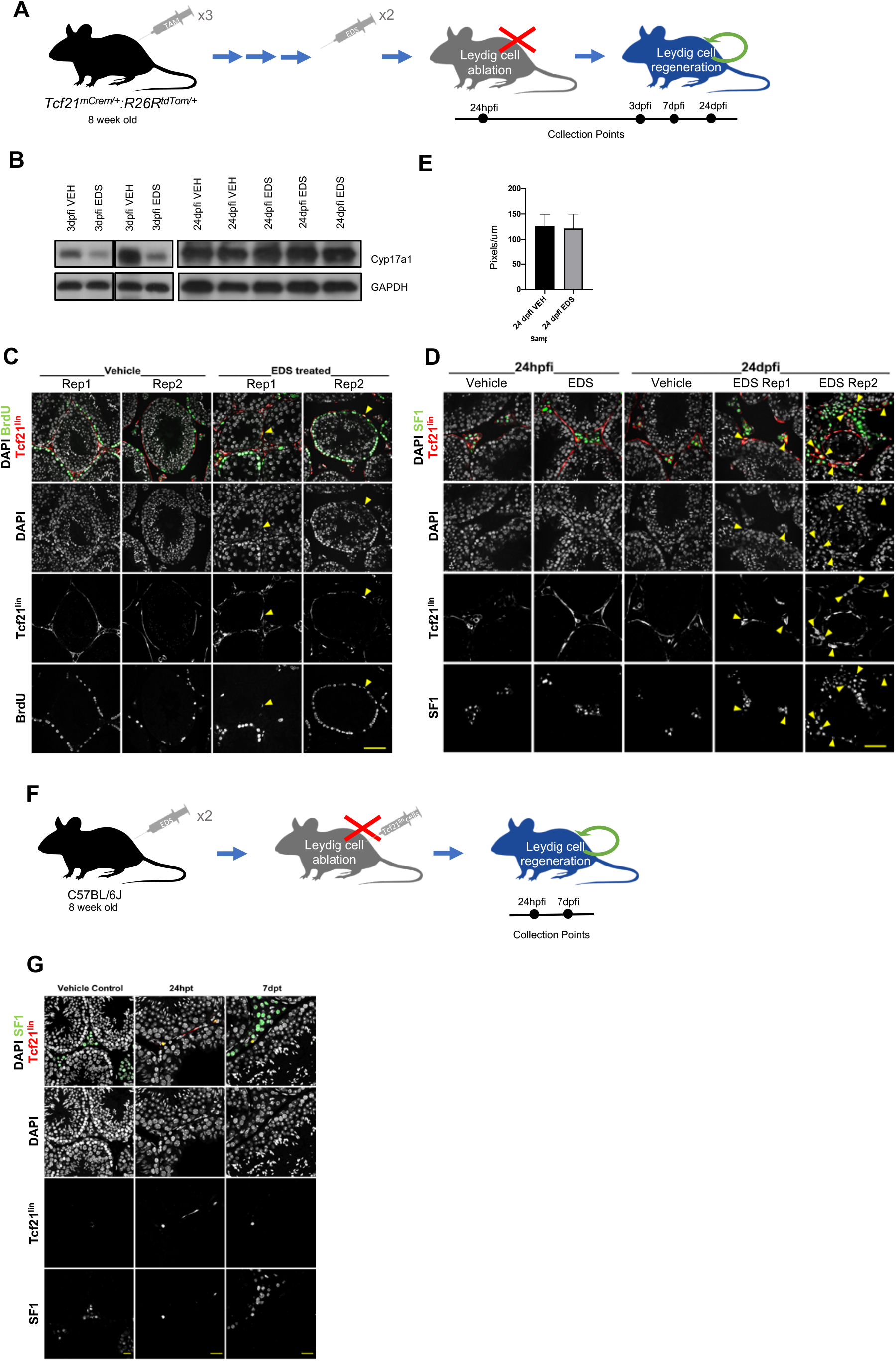
The Tcf21^lin^ population regenerates Leydig cells in vivo. **(A)** Schematic representation of the chemical method used to ablate Leydig cells in Tcf21^lin^ background. **(B)** Western blot illustrating Cyp17a1 protein decrease 3dpfi in EDS treated animals, and recovery of expression by 24dpfi; (**C)** Representative images of actively dividing Tcf21^+^ cells at 3dpfi, demarcated by TdTom/BrdU overlap. (**D)** Representative images of Leydig cells after regeneration illustrating co-expression of SF1/tdTom within EDS or vehicle injected *Tcf21^mCrem^:R26R^tdTom^* animals at 24dpfi. **(E)** Average Leydig cell diameter measurements in *Tcf21^mCrem^:R26R^tdTom^* EDS or vehicle injected animals, after Leydig cell recovery at 24dpfi (n=3, n=3). **(F)** Schematic representation of transplantation of Tcf21^lin^ cells following chemical ablation of Leydig cells in C57BL/6 animals. (**G)** Representative images of EDS treated C57BL/6 testes collected 24 hours and 7 days after transplant of Sca1^+^/cKit^-^ cells from *Tcf21^mCrem^:R26R^tdTom^* animals, co-immunostained with SF1/TdTom. Scale bars: 20 μm.

We next sorted adult Sca1^+^ cells from *Tcf21^mCrem^:R26R^tdTom^* animals and transplanted them into the testis interstitium of EDS-treated C57BL/6 animals (**Figure 5F**). We reasoned that if the transplanted Tcf21^lin^ population contributed to Leydig cell regeneration, then we should detect both Tcf21^lin^ and SF1 double positive cells in the C57BL/6 EDS-treated animal. We were encouraged to find that by 24hpfi, Tcf21^lin^ cells homed to the basement membrane, and that by 7dpfi, we began detecting Tcf21^lin^ and SF1^+^ cells (**Figure 5G**), suggesting that the Tcf21^lin^ cells engrafted in the ablated C57BL/6 mouse testis and gave rise to SF1^+^ cells (**Figure 5G**).

It is known from previous studies that Sertoli cells can be replaced by transplantation but cannot be regenerated following DTX treatment (Rebourcet et al., 2014a; Rebourcet et al., 2014b; Yokonishi et al., 2020). Our work demonstrates that Leydig cells can be regenerated from adult Tcf21^+^ progenitors, but whether this regenerative capacity extends to other somatic populations is not clear. To answer these questions for the peritubular myoid cells, we treated MYH11^cre-egfp^; Rosa26^iDTR/+^ mice (referred to as MYH11-cre:iDTR hereafter) with multiple low doses of DTX to ensure both animal survival and myoid cell ablation, and collected testes at 12hpfi and 4dpfi (**Figure S4A**). By 12hpfi, TUNEL positive cells were present on the tubule basement membrane in MYH11^cre-egfp^; Rosa26^iDTR/+^ mice but absent in control mice (**Figure S4B)** and no longer detectable by 4dpfi. However, the effect of DTX treatment - presence of vacuoles and disordered tubules - was visually persistent in hematoxylin and eosin stained cross-sections (**Figure S4B,C**). By 4dpfi, we also find a significant increase in BrdU positive cells surrounding the basement membrane (**Figure S4D**). Some of these are SMA^+^ whereas others are SMA^-^ (**Figure S4D)**, indicating that neighboring peritubular smooth muscle cells or interstitial cell progenitors re-enter the cell cycle to repair or regenerate the basement membrane (**Figure S4D,E**).

Since there is no known chemical approach that selectively eliminates peritubular cells in the testis, we therefore could not track peritubular myoid cell regeneration in *Tcf21^mCrem^:R26R^tdTom^* animals. As an alternate strategy, we transplanted adult Sca1^+^ interstitial cells isolated from *Tcf21^mCrem^:R26R^tdTom^* males into adult *MYH11^cre-egfp^; Rosa26^iDTR/+^* animals (**Figure S5A**). Although expression of MYH11^cre-egfp^ in organs outside of the testes proved to be challenging with this approach (most animals died 24 hours post-transplant), we were able to detect adult spindle-like Tcf21^lin^ cells at 24hpt surrounding the damaged tubules but we failed to detect any adult Tcf21^lin^ cells that had begun to express SMA at this time point (**Figure S5B**). In short, we have demonstrated that peritubular myoid cells can be regenerated and posit that the regeneration may rely on neighboring smooth muscle or interstitial progenitor. Whether this is a Tcf21^lin^ cell could not be ascertained.

### Adult Tcf21+ cells contribute to somatic turnover in the testis during aging

Once established, the somatic cells of the testis are maintained throughout a male’s reproductive age. However, a study in rats using [3H]thymidine labeling to detect proliferation suggested that Leydig and possibly peritubular myoid cells may undergo rare events of cellular turnover during an animal’s lifespan (Teerds et al., 1989). Consistent with this report, our scRNA-seq data in the adult mouse testis suggests that a rare subset of adult Tcf21^lin^ cells coexpress Cyp17A1 or Myh11 (data not shown), hence continued differentiation of Tcf21^+^ cells is possible, albeit quite rare and would not have been noticed if not for single-cell analyses. We asked whether adult mouse testis somatic cells might turnover during natural aging; and if so, if new cells would be derived from Tcf21^lin^ progenitors. To answer this question, we performed an extended lineage tracing experiment where five-week-old (adolescent) and 8-week-old (adult) *Tcf21^mCrem^:R26R^tdTom^* animals were injected with a single dose of 2 mg tamoxifen. Animals were euthanized either one week past final injection or 1 year past final injection (aged mice) (**Figure 6A**). In animals with short-term labeling we detected a few rare Tcf21^lin^ cells surrounding the basement membrane, and these cells do not overlap with Leydig cell markers such as SF1 (**Figure 6B)**. In the aged mice, we detected both Tcf21^lin^ and SF1^+^ cells as well as more labeling around peritubular cells suggesting that peritubular cells are also replenished (**Figure 6B**), and this labeling pattern varied slightly by individual animal which can be due to tamoxifen injection efficiency or true natural biological variability in inbred mice. Furthermore, similar results were also observed when using adolescent mice (5 weeks) (**Figure S6A,B**), although there was perhaps slightly broader labeling in the testis. Taken together, these data indicate that somatic cells of the testis naturally turnover at very low rates, and the Tcf21^lin^ population serves to replenish and maintain the adult Leydig cell population, and likely peritubular myoid cells as well.

**Figure 6:**
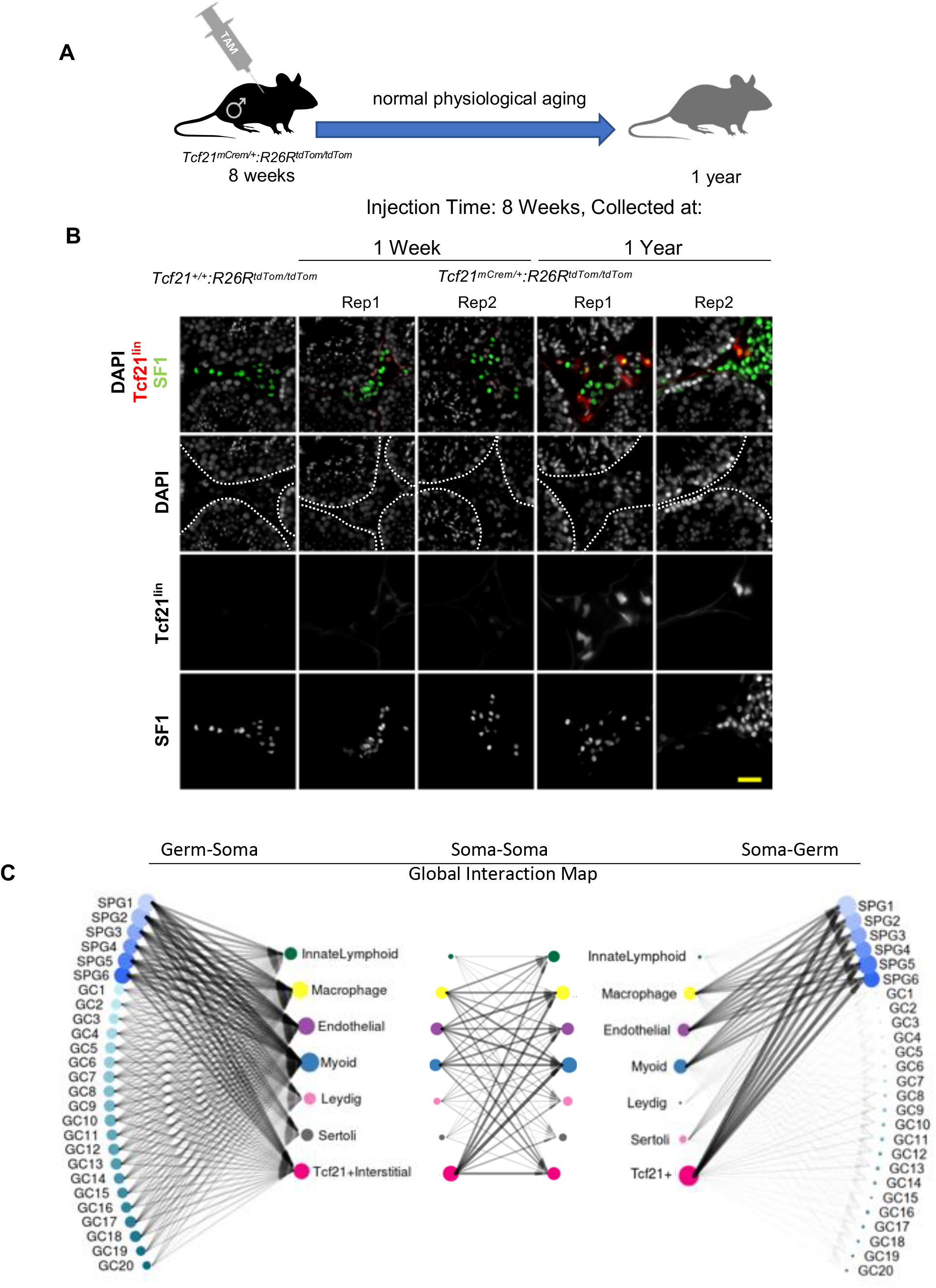
The Tcf21^lin^ population maintains testis tissue homeostasis during aging. **(A)** Schematic representation of Tcf21 lineage tracing in the aging testis. **(B)** Colocalization of Tcf21^lin^ (tdTomato), Leydig cell marker SF1 (green) and the nuclear counterstain DAPI (white) at one week or one year following injection of 8 week-old adults. Dotted lines indicate seminiferous tubules. Scale bars: 20 μm. **(C)** Summary of putative ligand – receptor interactions in the mouse testis between the germline and soma (left), within soma (center), and soma and germline (right). Arrows summarize top 5% of all interactions. scRNAseq data from Green et al., 2018, processed as described in Shami et. al., 2020. Symbol size indicates the number of receptor-ligand interactions contributed by a cell type and line width shows the number of interactions between the two cell types.

To get a sense of cellular interactions of Tcf21^+^ cells within the testis, we calculated Interaction Scores for highly variable and previously defined ligand-receptor (L-R) pairs in germ cells and somatic cells or among somatic cells of the testis (**Figure 6C, Table S3**). We previously showed that the Tcf21^+^ population has more potential interactions with spermatogonial populations than other germ cells (Shami et al., 2020) (**Figure 6C,** right panel), Similarly, we find that spermatogonia can convey information back to the Tcf21^+^ population (**Figure 6C**, left panel) via Pdgfa, various ADAMs, calmodulins, FGFs, and guanine nucleotide binding proteins (**Table S3**).

Within the somatic compartment, Tcf21^+^ cells have the greatest potential interactions with endothelial, myoid, and macrophages (**Figure 6C,** middle panel) including receptor-ligand signaling systems including Lrp1 (Cd91), a multifunctional, endocytic receptor capable of binding a vast array of ligands (May and Herz, 2003) and known to regulate the levels of signaling molecules by endocytosis, as well as directly participate in signaling for cell migration, proliferation, and vascular permeability (reviewed in (Lillis et al., 2005)). Like other tissue mesenchymal progenitors, Tcf21^+^ cells may also modulate local inflammatory responses, as they express Thrombomodulin (Tbhd) which interacts with and can proteolytically cleave the pro-inflammatory molecule Hmgb1 (reviewed in (Li et al., 2012)). Additionally, there are several more cell-specific interactions with myoid cells (Tgfb2-Tgfbr3; Gpc3-Cd81), endothelial cells (Cxcl12-Itgb1; Pdgfa-Pdgfra; and Vegfa-Itgb1), and macrophages (Igf1-Igf1r; F13a1-Itgb1) (**Table S3**). Several of these putative interactions are involved in growth, wound healing, phagocytosis, and matrix remodeling in various mesenchymal cell types (Mitchell and Mutch, 2019).

### The Tcf21^+^ population in the testis resembles resident fibroblast populations in other tissues

Finally, given the role of Tcf21^+^ cells in mesenchymal development of other tissues, including the heart, lung, and kidney (Acharya et al., 2011; Lu et al., 2000; Quaggin et al., 1999; Robb et al., 1998), we sought to understand whether similarities exist between the adult testis Tcf21 expressing population and other Tcf21^+^ mesenchymal populations found in single-cell analyses of other tissues. Towards this goal, we compared the transcriptome profile of our testis Tcf21^+^ cells with those from publicly available scRNA-seq datasets from coronary artery, heart, lung, and liver (Dobie et al., 2019; Farbehi et al., 2019; Wirka et al., 2019; Xie et al., 2018; Xiong et al., 2019). Among the cell types profiled in other organs, the testis Tcf21^+^ population most closely resembled the resident fibroblast or myofibroblast populations (**Figure 7, left**), as well as the transient fibroblast/myofibroblast populations that appear transiently after tissue injury (**Figure 7, right**). While diverse fibroblast cell types have been documented across tissues and may differ with respect to cellular markers or nomenclature (reviewed in (Swonger et al., 2016)), our results indicate that many of the fibroblast or myoblast populations some which are already implicated in fibrotic damage and tissue regeneration, are transcriptionally similar to the testicular Tcf21^+^ population characterized here, and may be playing similar roles in regulating responses to damage.

**Figure 7:**
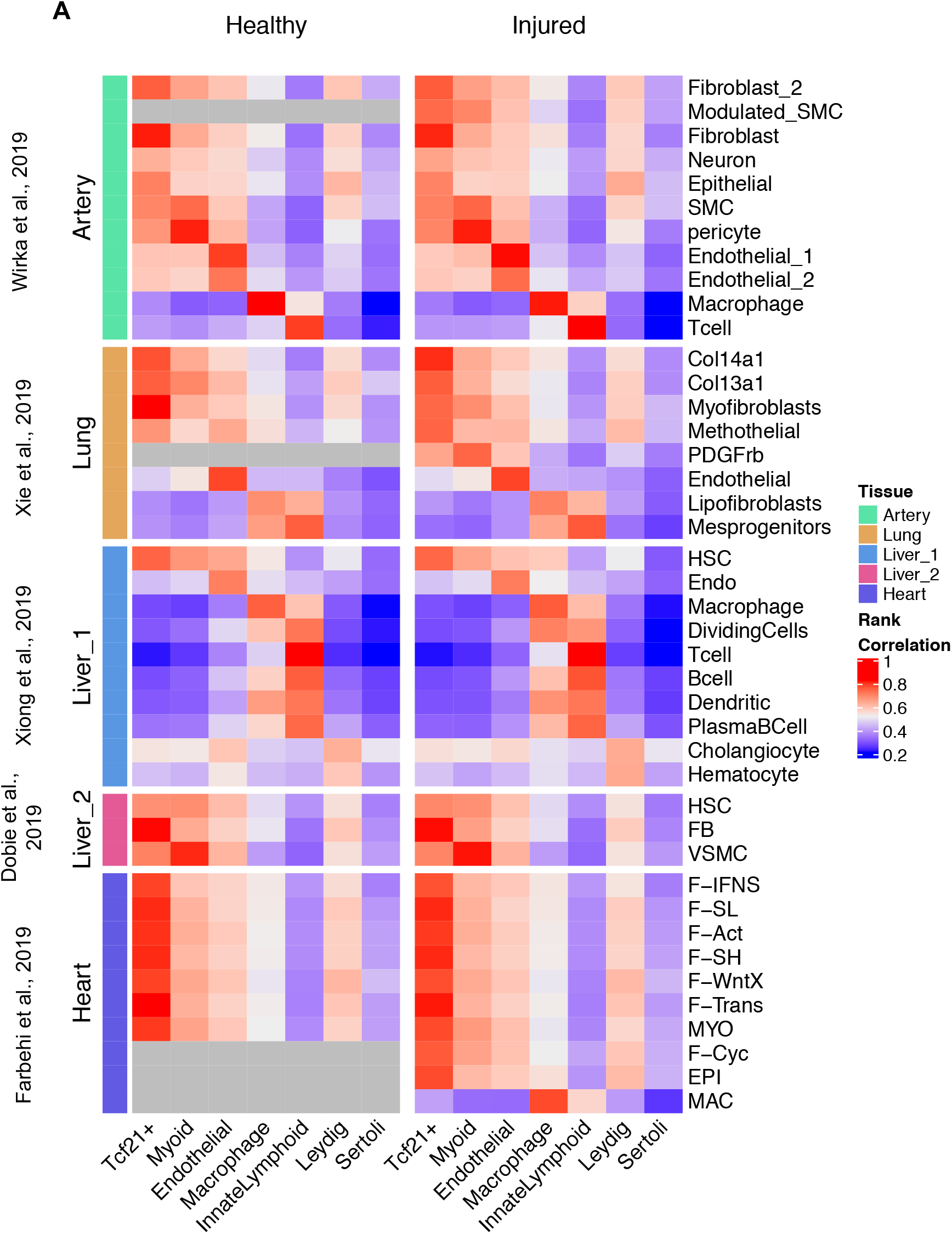
The adult Tcf21^+^ interstitial population resembles fibroblasts in other tissues. **(A)** Rank correlation of scRNA-seq cluster centroids of somatic cells from wild type adult mouse testes with other tissues in healthy adult mice (left panels) or after injury (right panels). Grey blocks indicate cell types not present in healthy datasets.

## Discussion

In this study, we sought to characterize functional properties of a population of Tcf21 expressing interstitial cells that we and others discovered in adult rodents by scRNA-seq (Green et al., 2018; Jung et al., 2019). We provide multiple lines of evidence to support that the Tcf21^+^ population has mesenchymal stem cell properties and is capable of producing multiple somatic cell types *in vitro and* is a bipotential somatic progenitor that contributes to multiple somatic lineages in the male and female gonads *in vivo*. In the adult testis, the Tcf21^+^ populations serve as a resident somatic stem cell that replenishes somatic cells during normal turnover, and responds to acute damages.

Tcf21 is a basic helix-loop-helix transcription factor, known to have roles in the development of numerous organs, including the testis (Lu et al., 2000; Lu et al., 2002; Quaggin et al., 1999). During testis development, Tcf21 is expressed in the bipotential gonadal ridge at E10.5 similar to other key transcription factors, including WT1 and Gata4 (Barsoum et al., 2009). Loss of Tcf21 resulted in gonadal dysgenesis (Cui et al., 2004; Quaggin et al., 1999). However, the fate of Tcf21 cells in the testis is not known. Our lineage tracing data show that the fetal Tcf21 + population is a bipotential gonadal progenitor giving rise to most somatic cell types including steroid-producing cells (fetal and adult), stromal/interstitial cells, and supporting cells, as well as vasculature, consistent with a common bipotential progenitor model recently described (Stevant et al., 2019; Stevant et al., 2018). Gonadal organogenesis is a complex process with multiple somatic progenitors have been described in both males and females. In males, WT1^+^ progenitors at E10.5 give rise to Sertoli cells, fetal Leydig and interstitial cells, and adult Leydig cells (Liu et al., 2016). Additionally, others have shown that CoupTFII, Nestin, Pdgfra positive cells contribute to testis cord, interstitial, and /or vasculature (Brennan et al., 2003). To a certain extent, our data reconcile these findings by showing that Tcf21 overlaps with many of these markers, but these populations appear to be molecularly and cellularly heterogeneous.

Furthermore, we find that Tcf21-expressing cells share characteristics with MSCs, adult stem cells that can self-renew *in vitro* and possess numerous traits like homing ability, multi-lineage potential, and secretion of molecules with anti-inflammatory and immunoregulatory effects (Dominici et al., 2006; Nauta and Fibbe, 2007). Like MSCs, adult Tcf21^+^ cells can be expanded *in vitro*, are highly clonogenic, and differentiate into osteogenic, chondrogenic, and adipogenic lineages. Additionally, we show that the Tcf21^+^ cells *in vitro* could be directly differentiated into smooth muscle and steroid-producing Leydig cells. Our scRNA-seq reveal that *in vitro* adult Leydig cell differentiation involves transition through a transient myofibroblast-like state that produces extracellular matrix components possibly supporting continued differentiation – which differs from the current model of Leydig cell differentiation in vivo. Interestingly though a few previous studies have detected smooth muscle actin and desmin in stem Leydig cells (SLCs) (Davidoff et al., 2004; Landreh et al., 2013; Stanley et al., 2012) but it was assumed that these gene products were contaminants from neighboring cell types. Although our in vitro-derived Leydig cells do not perfectly resemble or follow previously described trajectories in vivo, we successfully induced formation of mature steroid-producing cells in culture, findings that may be applicable to future *in vitro* and *in vivo* studies of Leydig cell development and testosterone production.

Testosterone is essential for the development and maintenance of male characteristics and fertility. Reduced serum testosterone affects millions of men and is associated with numerous pathologies including infertility, cardiovascular diseases, metabolic syndrome, and decreased sexual function. Although exogenous replacement therapies are largely successful in ameliorating these symptoms, they carry increased risks of cardiovascular and prostate disease or infertility (Baburski et al., 2019; Patel et al., 2019; Yeap et al., 2018). Thus, there is a need to develop approaches for transplanting testosterone-producing cells or activating endogenous progenitors to produce testosterone. The population described here could potentially serve as a source for future studies of cell-based testosterone replacement, as stem Leydig cells have the potential to provide a more natural means of androgen replacement to stimulate reproductive function in hypogonadal males and combat age-related decline in testosterone levels (Ge et al., 2005; Stanley et al., 2012). In support of this possibility, we show that allogenic Tcf21+ cell transplants or resident Tcf21+ within the testis can be activated to support tissue regeneration and replenish somatic cells in the aging testis. To date there have been several proposed putative stem Leydig cell populations have been described in fetal and early postnatal testis, which are demarcated by expression of diverse cell surface markers that are often species specific, like CD90 for rat and CD51 for mouse (reviewed in (Chen et al., 2017; Chen et al., 2020; Stanley et al., 2012; Ye et al., 2017). Furthermore, recent data suggests that these stem cell exhibit species specific properties. For example, Pdgfra^+^ cells isolated from rat can be differentiated to Leydig cells, whereas, although the PDGFRA+ isolated cells from the human testes exhibit aspects of MSC characteristics *in vitro*, but they are unable to fully differentiate into Leydig cells, nor can they produce testosterone (Eliveld et al., 2019).

Finally, given the critical role of Tcf21 in the development of other tissues, as well as in aging and disease models (Kikuchi et al., 2011; Maezawa et al., 2014), our findings for the testis revealed commonalities with other organ systems and disease states. For example, recent studies of atherosclerosis show that Tcf21-expressing cells are critical for the process of “phenotypic modulation”, where smooth muscle cells (SMCs) of the vasculature undergo a dedifferentiation process to form the atherosclerotic lesions. Specifically, the authors show that resident SMCs upregulate Tcf21 to transform into a fibroblast-like phenotype *in vivo* in mouse models of atherosclerosis, while loss of Tcf21 inhibits this process (Wirka et al., 2019). During development, a reverse pattern can be observed where Tcf21 is downregulated in cells fated to become mature coronary SMCs, while expression remains in fibroblasts (Acharya et al., 2011), This is similar to what we observe in vitro, where Tcf21 progenitors form fibroblast-like intermediates during the differentiation to Leydig cells. Thus, in both homeostatic and disease states, varying Tcf21 expression appears to play a role in modulating cell fate decisions in response to the local environment. Additionally, the adult testis Tcf21 populations have close transcriptional similarities with fibroblast or fibroblast-like populations in lung and liver, with similar functional roles in normal tissue maintenance and response to injury or disease. Since the response to tissue injury is often context dependent, it is important to understand what signaling pathways may influence key decisions such as fibrosis vs. regeneration. We show that in the testis Tcf21 has regenerative potential; while other have observed fibrosis in the testes of men with impaired spermatogenesis, and fibrosis is historically regarded as a hallmark of infertility (Adashi et al., 1996; Frungieri et al., 2002a; Frungieri et al., 2002b; Mayer et al., 2016; Meineke et al., 2000). However, whether dysregulation of the Tcf21^+^ population may be involved in the pathogenesis of infertility remains to be examined. A greater understanding of this population and its regulation may contribute to more informed strategies to restore testis function, as well as the tissue regeneration and tissue repair therapies for other organs. Furthermore, it will be important to determine whether Tcf21^+^ cells from one organ can be used to repair cells in alternative organs, or if the Tcf21^+^ populations across organs share a common origin.

In summary, we identify a multipotent interstitial Tcf21^+^ population in the mouse testis which contributes to gonadal somatic lineages, testis tissue homeostasis, and aging, and functions as a cellular source for Leydig and myoid cell regeneration following damage in adult mice. We further demonstrate methods for enrichment, differentiation, and modulation of adult Tcf21^+^ cells, with potential as a platform for studying testicular development, pathology, and novel therapeutics (Li et al., 2019).

## Acknowledgements

We thank members of the Hammoud, Li and Yamashita Labs for scientific discussions and manuscript comments. We thank Drs. Barry Zirkin and Haolin Chen for EDS compound and comments on manuscript. This research was supported by National Institute of Health (NIH) grants 1R21HD090371-01A1 (S.S.H., J.Z.L.), 1DP2HD091949-01 (S.S.H.), R01 HD092084 (K.E.O., S.S.H), F30HD097961 (A.N.S), F31HD100124 (G.L.M.), training grants 5T32HD079342 (A.N.S.), 5T32GM007863 (A.N.S.), NSF 1256260 DGE (L.M.), Rackham Predoctoral Fellowship (L.M.), T32GM007315 (L.M), T32HD007505 (G.L.M.), T32GM007315 (G.L.M.), and Michigan Institute for Data Science (MIDAS) grant for Health Sciences Challenge Award (J.Z.L., S.S.H.), Open Philanthropy Grant 2019-199327 (5384) (S.S.H.).

## Author contributions

S.S.H., H.L., and A.N.S. provided overall project design. Y.S., H.L., A.N.S., L.M., G.L.M., M.S., M.C., C.S., and J.C. performed experiments. Q.M. J.Z.L, and X.Z. analyzed data. A.N.S and S.S.H. wrote the manuscript with input from H.C., J.S.R., K.E.O, M.T. and J.Z.L. Comments from all authors were provided.

## STAR Methods

### LEAD CONTACT AND MATERIALS AVAILABILITY

Further information and requests for resources and reagents should be directed to and will be fulfilled by the Lead Contact, Saher Sue Hammoud.

### EXPERIMENTAL MODEL AND SUBJECT DETAILS

#### Mice

All animal experiments were carried out with prior approval of the University of Michigan Institutional Committee on Use and Care of Animals (Animal Protocols: PRO06792, PRO00006047, PRO00008135), in accordance with the guidelines established by the National Research Council Guide for the Care and Use of Laboratory Animals. Mice were housed in the University of Michigan animal facility, in an environment controlled for light (12 hours on/off) and temperature (21 to 23°C) with ad libitum access to water and food (Lab Diet #5008 for breeding pairs, #5LOD for non-breeding animals).

Colony founders for Rosa26^iDTR^(Stock #007900), Cyp17a1-iCre(Stock #028547) and MYH11^cre-Egfp^ (Stock #007742)(maintained on a B6 background), Oct4-Egfp (Stock #004654), *PDGFRA^EGFP^*(Stock007669), and *R26R^tdTom^* mice (Stock #007909) were obtained from Jackson Labs. The *Tcf21^mCrem^ and the Gli1^EGFP^* were generously provided by Michelle Tallquist and Deb Gumuccio, respectively. All EDS injection studies were performed on age matched C57BL/6 mice obtained from Jackson Labs (Stock #000664).

For detailed mouse strain information, see below and Key Resources Table.

### METHOD DETAILS

#### Interstitial Populations Single Cell Data Analysis

##### Single cell RNA-sequencing analysis comparing somatic cell types in the adult mouse testis

Somatic cells (N=3,622) and their cell type classifications were previously described by (Green et al., 2018). To compare somatic cell populations of the testis we obtained the Euclidean distances for the somatic cell centroids then ordered the cell types using the optimal leaf ordering (OLO) algorithm in R Package Seriation. Based on this analysis, we discovered that the Tcf21 population is highly correlated to endothelial, myoid and Leydig cells, but very distinct from immune cells and Sertoli cells. To get a better understanding of the functional role of the Tcf21 population, we called differentially expressed genes in each somatic cell type using a nonparametric binomial test. The differentially expressed genes have: (1) At least 20% difference in detection rate; (2) a minimum of 2-fold change in average expression levels, and (3) p-value < 0.01 in the binomial test. Pathway Enrichment analysis for the differentially expressed genes was performed with PANTHER tool v.15 (http://www.pantherdb.org) (Mi et al., 2019). Significance of the over- and underrepresentation of GO Complete Biological Process categories was calculated using Fisher’s exact test and multiple testing correction with the false discovery rate.

##### Ligand-receptor analysis

We used a previously curated list of ligand-receptor pairs in human as reference, identifying corresponding orthologues in mouse as previously described (Shami et al., 2020). We selected pairs with either highly variable ligand gene among the 7 somatic cell centroids, or highly variable receptor gene among the 26 germ cell centroids (6 SPG and 20 non-SPG clusters). Thresholds were set for both genes mean and variance across the clusters according to the density. For each ligand-receptor pair we calculated its apparent signaling strength as an “Interaction Score”, defined as the product of the mean expression level of the ligand in one cell type and that of the receptor in another cell type. In all, we calculated such an Interaction Score matrix of cell type pairs for germ(ligand)-soma(receptor) interaction, soma (ligand)-soma (receptor) interaction and soma (ligand)-germ(receptor) interaction (reproduced with permission from (Shami et al., 2020), respectively. To extract the general signaling pattern for each interaction matrix, we defined “strong interactions” for each matrix by keeping the highest 5% Interaction Scores for each matrix. We then calculated the number of such strong L-R interactions for each pair of cell types as their overall interaction strength, and displayed them as the line width of arrows in the pairwise interaction plots.

##### Cell type correlations across tissues

We downloaded the single cell counts data from GEO for artery, lung, heart and two liver datasets (Dobie et al., 2019; Farbehi et al., 2019; Wirka et al., 2019; Xie et al., 2018; Xiong et al., 2019). For the lung, heart datasets, and the liver dataset from Xiong et al, we downloaded the accompanied cell type labels for single cells. For each cell type in each tissue, including the testis cell types, we calculated its expression centroid as the mean of the normalized gene expression profile of all cells belonging to that cell type. We then calculated the spearman rank correlation for all cell type pairs between testis and other tissues. For the artery dataset and the liver dataset from Dobie et al, the cell type labels for single cells are not available. We analyzed the raw data using Seurat by following the analysis descriptions from their original papers. For parameters that are not specified, we either used default values or set accordingly. Finally, we clustered the single cells by Louvain clustering in Seurat to get clusters comparable to the results in the papers and assigned cell types according to gene markers provided in the two papers. We then followed the same procedure to calculate cell type expression centroid and spearman rank correlations with cell types from testis. All the datasets included healthy and injured samples which were processed separately in calculations.

#### In Vitro Differentiation Assays

##### Flow cytometry

Testes were collected from adult C57BL/6 (JAX^®^mice, stock #000664) mice and enzymatically and mechanically dissociated into a single cell suspension. Briefly, testes from adult mice were excised and the tunica albuginea was removed. Seminiferous tubules were transferred to 10ml of digestion buffer 1 (comprised of Advanced DMEM:F12 media (ThermoFisher Scientific), 200 mg/ml Collagenase IA (Sigma), and 400 units/ml DNase I (Worthington Biochemical Corp). Tubules were dispersed by gently shaking by hand and a five-minute dissociation at 35°C/215 rpm. To enrich for interstitial cells, tubules were allowed to settle, and the supernatant was collected, quenched with the addition of fetal bovine serum (FBS) (ThermoFisher Scientific), filtered through a 100 um strainer and pelleted at 600g for 5 minutes. The remaining tubules were then transferred to digestion buffer 2 (200 mg/ml trypsin (ThermoFisher Scientific) and 400 units/ml DNase I (Worthington Biochemical Corp) dissolved in Advanced DMEM:F12 media) and dissociated at 35°C / 215 rpm for 5 min each and quenched with the addition of fetal bovine serum (FBS) (ThermoFisher Scientific). The cell pellets from multiple digests were combined and filtered through a 100um strainer, washed in Phosphate-buffered saline (PBS), pelleted at 600g for 3min, and re-suspended in MACS buffer containing 0.5% BSA (MiltenyiBiotec).

The single cell suspensions were stained with a single antibody or combination of antibodies depending on the experiment. The antibodies used include anti-Ly6a-Alexa Fluor 488 (1:100; Biolegend, Cat#108115), Biotinylated anti-Ly6a (1:200; Biolegend, Cat#108103), streptavidin conjugated Alexa Fluor 488 (1:1000; Life Technologies Cat# S11223; RRID: AB_2336881), anti-CD73-APC (1:300; Biolegend, Cat#127209), anti-CD90.1-Brilliant Violet 650 (1:300; Biolegend, Cat#202533), anti-CD29-PE/Dazzle 594(1:300; Biolegend, Cat#102231), anti-CD105-PerCP/Cy5.5 (1:300; Biolegend, Cat#120415), anti-CD45-Brilliant Violet 510 (1:300; Biolegend, Cat#), anti-CD117-PE/Cy7 (c-KIT) (1:300; Biolegend, Cat#105813), and anti-CD34-PE (1:300; Biolegend, Cat#128609).

##### Tri-lineage differentiation assay

Testes were collected from adult C57BL/6 or *Tcf21^mCrem^:R26R^tdTom^* mice and dissociated into a single cell suspension and sorted for Sca1^+^/cKit^-^ or Sca1^+^/ *Tcf21^lin^*/cKit^-^cells, respectively. For adipogenic differentiation, 3 × 10^4^ cells were plated in a monolayer, cultured for 10 days (StemProAdipogenic differentiation kit) and stained with either Oil Red O or Perilipin. For chondrogenic differentiation, 5 × 10^4^ cells were plated in micromass, cultured for 21 days (StemProChondrogenic differentiation kit) and stained with either Alcian blue or Sox9. For osteogenic differentiation, 1 × 10^4^ cells were plated in micromass, cultured for 14 days (StemPro Osteogenic differentiation kit), and stained with either Alizarin red or Osterix (Rux et al., 2016).

##### CFU-F assay

Colony forming unit assays were performed as previously described (Park et al., 2012). Briefly, testes were collected from C57BL/6 or adult *Tcf21^mCrem^:R26R^tdTom^* males following 3 injections of tamoxifen (2 mg) every other day. Following single cell dissociations, Sca1^+^/cKit^-^, Sca1^+^/*Tcf21^lin^*/cKit^-^, *Tcf21^lin^*/cKit^-^, or cKit^+^/Sca1^-^ cells were plated into Corning Primaria 6 well plates at a density of 1000 cells/well. Cells were cultured for 14 days in MesenCult MSC medium (StemCell Technologies) and colonies were stained using Giemsa. Colonies were defined as clumps having either >20 or >50 cells.

##### In vitro directed differentiation to Leydig and myoid cells

Testes were collected from adult C57BL/6 or *Tcf21^mCrem^:R26R^tdTom^ males*, dissociated into a single cell suspension, and sorted for Sca1^+^/cKit^-^ cells. Cells were plated into a 24 well, Matrigel-coated plates at a density of 100,000 cells/well in DMEM/F12 supplemented with 10% FBS and 1X Normocin. For myoid cell differentiation, after 18 hours media was replaced with differentiation media-DMEM/F12 supplemented with 1X Penicillin/streptomycin, 10 ng/ml PdgfAA, 10 ng/ml PdgfBB, 0.5 uM SAG, 10 ng/ml BMP2, 10 ng/ml BMP4, 10 ng/ml Activin A, and 1 mM Valproic acid. After the 7-day differentiation protocol, cells were stained for SMA (1:250, Sigma). For Leydig cell differentiation, the FACs sorted cells were initially recovered for 18 hours in DMEM+10%FBS and then the cells were expanded for 3 days in DMEM/F12 supplemented with 1X Normocin, 10 ng/ml PdgfAA, 10 ng/ml PdgfBB, 0.5 uM SAG, and 10 ng/ml Fgf2. After 3 days, the expansion media was replaced with differentiation media-DMEM/F12 supplemented with 1X Penicillin/streptomycin, 10 ng/ml PdgfAA, 10 ng/ml PdgfBB, 0.5 uM SAG, 10 ng/ml Fgf2, 5 mM LiCl_2_, and 10 uM DAPT. After 10 days in differentiation media, cells were stained for SF1 (1:100, Cosmo Bio). Media was collected every other day for testosterone measurements.

###### Clonal expansion and directed differentiation of Tcf21^+TdTom+^ cells

To assess multipotency of *Tcf21^lin^* cells, testes were collected from adult *Tcf21^mCrem^:R26R^tdTom^* males and dissociated into a single cell suspension. Individual Sca1^+^/*Tcf21^lin^*/cKit^-^ cells were sorted into Corning Primaria 96 well plates and cultured in MesenCult mouse MSC media for ~3 weeks to allow for colony formation. Individual colonies were then directed to differentiate to either a myoid or Leydig cell fate following the directed differentiation protocols described above. Cells were then stained for either SMA (for myoid cells) or SF1 (Leydig cells).

##### Drop-seq analysis of the Leydig cell differentiation time course analysis

Briefly, cells collected from various time points of differentiation were diluted to 280 cells/ul and processed as described previously (Macosko et al., 2015). Briefly, cells, barcoded micro particle beads (MACOSKO-2011-10, Lots 113015B and 090316, ChemGenes Corporation), and lysis buffer were co-flown into a microfluidic device and captured in nanoliter-sized droplets. After droplet collection and breakage, the beads were washed, and cDNA synthesis occurred on the bead using Maxima H-minus RT (Thermo Fisher Scientific) and the Template Switch Oligo. Excess oligos were removed by exonuclease I digestion. cDNA amplification was done for 15 cycles from pools of 2,000 beads using HotStart ReadyMix (Kapa Biosystems) and the SMART PCR primer. Individual PCRs were purified and pooled for library generation. A total of 600 pg of amplified cDNA was used for a Nextera XT library preparation (Illumina) with the New-P5-SMART PCR hybrid oligo, and a modified P7 Nextera oligo with 10 bp barcodes. Sequencing was performed on a NovaSeq (Illumina) for read 2 length of 94 nt with the Read1CustomSeqB primer. Oligo sequences are the same as previously described (Green et al., 2018; Macosko et al., 2015).

##### Single-cell RNA-seq Analysis Across Time points

The paired-end Drop-seq data from days 4, 7, and 14 of *in vitro* Leydig cell differentiation were sequenced in the same batch and were processed using *Drop-seq tools* v1.13 from the McCarroll laboratory as previously described (Macosko et al., 2015; Shekhar et al., 2016). Specifically, the reads were aligned to the mouse reference genome (GRCm38, version 38) using *STAR* v2.7.1a (Dobin et al., 2013). The pipeline generated digital gene expression matrices with genes as rows and cells as columns that served as the starting point for downstream analyses.

The cells from each time point were first filtered by cell size and integrity – cells with <500 detected genes or with >10% of transcripts corresponding to mitochondria-encoded genes were removed, resulting in 973 – 2980 pass-QC cells for the three time points, for a total of 6,124 cells. Among the retained cells, the average number of detected genes per cell was ~1,888, and the average number of UMIs was ~4,838. For each cell, we normalized transcript counts by (1) dividing by the total number of UMIs per cell and (2) multiplying by 10,000 to obtain a transcripts-per-10K measure, and then log-transformed it by E=ln(transcripts-per-10K+1).

For each time point, we standardized the expression level of each gene across cells by using (E-mean(E))/sd(E), and performed PCA using highly-variable genes (HVG). We obtained 4 clusters using Louvain-Jaccard clustering with top PCs by R package *Seurat* (v2.3.4). We calculated cluster centroids and ordered the clusters by minimizing pairwise Euclidean distance of cluster centroids in R package *Seriation*. We evaluated batch effect by comparing the top PC placements and rank correlation of ordered cluster centroids across the 3 time points. Differentially expressed markers for each cluster were obtained by comparing it against all other clusters using a nonparametric binomial test, requiring at least 20% higher detection rate, a minimum of 1.5-fold higher average expression level, and p-value < 0.01. Based on known markers, there is detected germ cell no in the data.

We extracted the somatic cells from the INT4 dataset from (Green et al., 2018). This dataset was enriched for the Tcf21^+^ interstitial population, and was used as the starting time point: day-0. We merged the 3 datasets of *in vitro* Leydig differentiation with the somatic cells of INT4, for 6,619 good-quality cells and 24,698 detected genes. These cells on average have 1,837 detected genes and 4,623 UMIs per cell. We selected 2,344 HVG genes in the merged dataset and did PCA using HVG. We performed t-SNE, UMAP and Louvain-Jaccard clustering using top PCs. We obtained 10 clusters initially, and ordered the clusters as described above. Based on differentially expressed markers for each cluster and rank correlation across the cluster centroids, we decided to merge 3 clusters (clusters 4-6) and identified them as ECM myofibroblast. We merged 2 other clusters (clusters 7-8) as differentiating myofibroblast. This led to the identification of 7 cell types for *in vitro* Leydig differentiation– 1. Endothelial, 2. Interstitial progenitor, 3. Proliferating progenitor, 4. ECM myofibroblast, 5. Differentiating Myofibroblast, 6. Differentiating Leydig, and 7. Leydig. We did pseudotemporal ordering of cells from the 4 time points (days 0, 4, 7, and 14) by *Monocle3* and visualized the single-cell trajectory in UMAP space.

We compared our *in vitro* Leydig differentiation data with those of fetal mouse gonads and adult mouse testis somatic cells. Specifically, we calculated the rank correlation between our 7 cell type centroids of Leydig differentiation data with the 6 cell type centroids from Nr5a1^+^GFP^+^ Progenitor cells from E10.5 – E16.5 fetal male mouse gonads (Stevant et al., 2018) using markers present in both data (N=2,692). We also calculated the rank correlation between our 7 cell type centroids with the 7 cell type centroids from adult mouse testis (Green et al. 2018) using the union of markers present in both datasets (N=1,769).

#### Tcf21 Lineage Tracing

##### Tcf21 lineage tracing analysis and co-localization with immunofluorescence in fetal and adult testis and ovary

*Tcf21^mCrem^:R26R^tdTom^* or *Tcf21^mCrem^:R26R^tdTom^;* Oct4-egfp timed-pregnant females were administrated with a single dose of 1 mg tamoxifen via gavage at E9.5, E10.5, E11.5 or E12.5. Embryos were obtained at E11.5, E17.5, or E19.5 via C-section. Tail clippings from the embryos were used to identify sex and genotype. Embryonic gonads were fixed in 4% paraformaldehyde (PFA) at 4°C for one hour, transferred to 30% sucrose in 1X PBS at 4°C overnight and embedded in OCT (Surgipathcryo-gel, Leica #39475237). To analyze the *Tcf21^lin^* contribution in adult testis and ovaries, the E19.5 pups obtained by C-section were fostered toCD1 females. The foster mice were euthanized at 10 weeks. Adult testes and ovaries were fixed in 4% PFA at 4°C overnight, transferred to 30% sucrose in 1X PBS at 4°C for overnight and embedded in OCT.

For immunofluorescence,7-10 micron thick OCT sections were cut using a Leica CM3050S cryostat and sections were refixed with 4% PFA for 10 minutes and permeabilized by incubation in 0.1% Triton in PBS for 15 minutes. Sections were blocked in 1X PBS, 3% BSA, and 500mM glycine for one hour at room temperature and co-stained with Fgf5, Sox9, 3β-HSD, CD34, SMA, Vasa, CoupTFII, CD31, SF1, WT1, Foxl2, and DsRed antibodies. The primary antibodies and concentrations used are summarized in the table below. All secondary antibodies (Alexa-488-, Alexa-568-, and Alexa-647-conjugated secondary antibodies; Life Technologies/ Molecular Probes) were all used at a 1:1000 dilution. DAPI was used as a nuclear counterstain at 1:1000. Representative images were taken with a Zeiss AX10 epifluorescence microscope, a Leica SP8 confocal microscope, or a Nikon A1R-HD25 confocal microscope and processed with ImageJ.

##### Quantification of immunofluorescence colocalization

Tissue sections were stained for immunofluorescence as described above and >20 images per testis were imaged with a 40X 1.2NA objective on a Zeiss AX10 epifluorescence microscope, all at a single z-section. The percent overlap between the *Tcf21^lin^* population and several marker proteins was done using a custom written ImageJ macro (available upon request). Briefly, nuclear regions of interest (ROIs) were created from DAPI staining by blurring with a Gaussian filter, making the image binary, separating overlapping nuclei with a watershed function, then saving the outline of each binary nucleus to ImageJ’s ROI manager. The signal from *Tcf21^lin^* and the immunostaining were made binary using ImageJ’s automatic thresholding function and the overlap of the binary stain and the nuclear ROI was measured using ImageJ’s Measurement function. Cells positive for each stain and double positive cells were sorted and identified in Microsoft Excel. The quantification from each testis was the sum of all quantified images taken from a single testis. At least six testes from at least three mice were imaged and quantified for each protein marker shown. The macro was optimized by contrasting its results to manual quantification from at least five images per immunostaining.

##### Lineage-tracing analysis of the Tcf21+ population in the aged testis

Five or Eight weeks old *Tcf21^mCrem^:R26R^tdTom^* males were injected with a single dose of 2 mg tamoxifen intraperitoneally (i.p.). The testis were collected one week (as control) or one year past injection. The testes were fixed in 4% paraformaldehyde at 4°C overnight, transferred to 30% sucrose in 1X PBS at 4°C overnight and embedded in OCT. The sections were co-stained for SF1.

#### In vivo ablation of Leydig and myoid cells

##### Ethane Dimethane Sulfonate (EDS) Injections

Ethane dimethane sulfonate (EDS) (AABlocks, Cat. No: 4672-49-5) was dissolved in DMSO at a concentration of 150 mg/mL. Working solutions of EDS were further diluted in PBS to a final concentration of 50mg/mL. A total of 300mg/kg of EDS was injected i.p. every other day for 2 days into C57BL/6 age-matched mice. Testes and sera were collected 12 hours, 24 hours, 4 days, 7 days, 14 days or 21 days past final injections. Negative controls were given vehicle treatment of DMSO: PBS without EDS.

##### Diphtheria Toxin Injections

Diphtheria toxin (DTX) (Sigma) was diluted in PBS at a concentration of 2 mg/ml. A final concentration of either 80ng or 150ngDTX was injected i.p. every day for 3 or 4 days into *Cyp17a1-icre;Rosa26^DTR^* or *Myh11^cre-egfp^;Rosa26^DTR^*, respectively. Testes and sera were collected 12 hours, 24 hours, 4 days, 7 days, 14 days or 21 days past final injections. Litter mate negative controls for either the Cre-recombinase, the diphtheria toxin receptor, or both, were given DTX and served as wild-type controls.

###### TUNEL staining and positive cell counts

Testes were collected, fixed in 4% PFA (in PBS) for approximately 16hrs at 4°C, dehydrated in ethanol wash series, and embedded in paraffin. Five-micron FFPE tissue sections were deparaffinized, rehydrated, and permeabilized in 20 μg/ml Proteinase K solution for 15 minutes at room temperature. Samples were further processed following the Promega DeadEnd Colorimetric TUNEL kit according to the manufacturer instructions. All images collected used a Leica Leitz DMRD microscope. The percent TUNEL positive cells was determined by averaging at least 100 tubules for each sample.

###### Hormone Measurements

Testosterone measurements were performed by the University of Virginia Center for Research in Reproduction Ligand Assay and Analysis Core. Blood was collected using a 1mL insulin syringe and left to coagulate at room-temperature for 90 minutes. Sera were collected as described on the Ligand Assay and Analysis Core Web site (http://www.medicine.virginia.edu/research/institutes-and-programs/crr/lab-facilities/sampleprocessingandstorage-page).

###### FFPE Immunofluorescence

Whole testes were fixed in 4% PFA overnight at 4°C and processed for formalin fixed paraffin embedding as described in (Fisher et al 2006). Five-micron FFPE tissue sections were deparaffinized by incubation in Histoclear 3x for 5 minutes, followed by incubation in 100% EtOH 2x for 5 minutes, 95% EtOH 2x for 5 minutes, 80% EtOH 1x for 5 minutes, 70% EtOH 1x for 5 minutes, 50% EtOH 1x for 5 minutes, 30% EtOH 1x for 5 minutes, and deionized water 2x for 3 minutes each. Tissue sections were permeabilized by incubation in 0.1% Triton in PBS for 15 minutes. For all antibodies, antigen retrieval was performed by boiling in 10mM sodium citrate, pH 6.0 for 30 minutes. Sections were blocked in 1x PBS, 3% BSA, 500mM glycine for three hours at room temperature. Endogenous peroxidases and alkaline phosphatases were blocked by a ten-minute incubation in BloxAll solution (Vector Labs, Cat. No: SP-6000). The primary antibodies and concentrations used are listed in table X. Alexa-488-, Alexa-555-, and Alexa-647-conjugated secondary antibodies (Life Technologies/ Molecular Probes) were all used at 1:000. DAPI was used as a nuclear counterstain.

###### Cell diameter measurements

For all cell diameter measurements, FFPE slides were stained as described above and costained with SF1to mark Leydig cells and 488-wheat germ agglutinin (WGA, Biotium Cat. No: 29022-1) to mark cell perimeter. A tubule was centered in the field of view at 40X magnification and an image was taken. Between 80-100 well-defined Leydig cells, marked by both SF1and clear visible cell perimeter by WGA, were counted per sample. Cell diameter was measured manually using ImageJ’s Line and Measure functions.

###### FFPE Immunohistochemistry

Whole testes were fixed in 4% PFA overnight at 4°C, processed and deparaffinized as described above. The primary antibodies and concentrations used are listed in table below. Horseradish peroxidase and alkaline phosphatase conjugated secondary antibodies (Abcam) were all used at a 1:100 concentration and left to incubate for 1-hour at room temperature. Slides were rinsed of developing solution under running DI water for 1 minute and mounted in Permount.

###### Transplanting the Tcf21^Lin^ cells into the testis of WT EDS treated animals or Myh11;DTR mice

6-18 week *Tcf21^mCrem^:R26R^tdTom^male* mice were injected with 1mg tamoxifen intraperitoneally every other day for a total of three injections. Testes were dissociated into a single cell suspension and the Sca1^+^;cKit^-^stained cells were collected by FACs, as described above. For each animal ~65,000-150,000 cells were diluted in 10ul of MEM media plus Trypan Blue and were injected into the interstitium via the rete testes of either EDS treated C57BL/6 animals 24hours after final injection (hpfi) of EDS or into *Myh11;DTR mice* 24 hpfi of DTX. As a control, 10uL of MEM media plus Trypan Blue was injected into the contralateral testis of each experimental animal. Testes were collected at 24 hours past transplant (hpt), 4 days past transplant (dpt) and 7dpt for FPPE processing as described above.

| **Antibody** | **Company** | **Product #** | **Application** | **Concentration** |
| --- | --- | --- | --- | --- |
| Ly6a-Alexa | Biolegend | 108115 | Flow cytometry | 1:200 |
| Fluor 488 |  |  |  |  |
| CD73-APC | Biolegend | 127209 | Flow cytometry | 1:300 |
| CD90.1-Brilliant Violet<br>650 | Biolegend | 202533 | Flow cytometry | 1:300 |
| CD29-PE/Dazzle 594 | Biolegend | 102231 | Flow cytometry | 1:300 |
| CD105-PerCP/Cy5.5 | Biolegend | 120415 | Flow cytometry | 1:300 |
| CD45-Brilliant<br>Violet 510 | Biolegend | 103137 | Flow cytometry | 1:300 |
| CD117-PE/Cy7<br>(c-KIT) | Biolegend | 105813 | Flow cytometry | 1:300 |
| CD34-PE | Biolegend | 128609 | Flow cytometry | 1:300 |
| Fgf5 | Santa Cruz | sc-376264 | immunofluorescence | 1:100 |
| Sox9 | EMD Millipore | ABE571 | immunofluorescence | 1:500 |
| Sox9 | Abcam | ab3697 | immunofluorescence | 1:500 |
| 3β-HSD | Santa Cruz | Sc-30820 | immunofluorescence | 1:200 |
| CD34 | Abcam | ab81289 | immunofluorescence | 1:200 |
| Smooth muscle<br>actin | Sigma | A5228 | immunofluorescence | 1:200 |
| Vasa (Ddx4) | Abcam | ab13840 | immunofluorescence | 1:200 |
| CoupTFII<br>(Nr2F2) | R&D Systems | PPH7147-00 | immunofluorescence | 1:300 |
| CD31 (PECAM) | BD Biosciences | 553370 | immunofluorescence | 1:100 |
| SF1 (Nr5A1) | Hammer lab |  | immunofluorescence | 1:500 |
| SF1 (Nr5A1) | Cosmo Bio | KAL-KO610 | immunofluorescence | 1:100 |
| WT1 | Santa Cruz | sc-192 | immunofluorescence | 1:100 |
| Foxl2 | Wilhelm lab |  | immunofluorescence | 1:300 |
| Foxl2 | Novus<br>Biologicals | NB100-1277 | immunofluorescence | 1:300 |
| DsRed (for<br>tdTomato) | Takara Bio USA<br>Inc | 632496 | immunofluorescence | 1:1000 |
| Cyp17A1 | ProteinTech | 14447-1-AP | Western | 1:500 |
| β-actin | Novus<br>Biologicals | NBP1-47423 | Western | 1:2000 |
| GAPDH | ProteinTech | 10494-1-AP | Western | 1:500? |
| BrdU | Abcam | Ab6326 | immunofluorescence | 1:500 |
| Osterix | Abcam | ab22552 | Immunofluorescence | 1:250 |
| Perilipin | Sigma |  | Immunofluorescence | 1:250 |

### DATA AND CODE AVAILABILITY

#### Data Resources

The accession number for the raw and processed data files are available in GEO:XXXXXX

## Supplemental Figure Legends

**Supplemental Figure 1: The Tcf21^+^ population is molecularly heterogeneous and expresses mesenchymal progenitor markers. (A)** Gating strategy used to characterize interstitial Sca1^+^ cells in the testis. The first plot shows side scatter area and CD45 fluorescent intensity used to eliminate immune cell populations (selection of Sca1^+^, CD45^-^). The second plot is a Sca1 and cKit fluorescent intensity plot used to remove Leydig cells (selection of Sca1^+^, CD45^-^, cKit^-^). All positive gates were determined using FMO isotype controls. **(B)** t-SNE analysis (perplexity 35,300 iterations) of thousands of cells co-immunostained for Sca1, CD73, CD105, Thy1, TdTomato, and CD34 identifies five Sca1^+^ subtypes based on the signal intensity of various markers. **(C)** Fluorescent intensity histogram of the 6 markers across the 5 clusters defined in panel B. **(D-G)** Immunofluorescence overlay of Tcf21^lin^ with PDGFRA^eGFP^ **(D)**, CoupTFII (**E**), CD34 **(F)** and Fgf5 **(G)** expression in the adult mouse testis. Scale bars: 20μm.

**Supplemental Figure 2: The Tcf21^lin^ population is a multipotent mesenchymal progenitor and can be differentiated to somatic cells of the testis. (A)** Schematic representation of experimental outlines for trilineage differentiation, CFU-F, and myoid/Leydig cell differentiation**. (B)** Representative images from Sca1^+^ trilineage differentiation. Adipogenic differentiation was analyzed at 10 days using Oil Red O, chondrogenic differentiation using Alcian blue at 14 days, and osteogenic differentiation at 21 days using Alizarin red. **(C)** Enrichment of clonogenic cells in the Tcf21^lin^ population. Left panels are CFU-F representative images and the right panel is the quantification of the number of colonies with either >20 or >50 cells. Data are presented as mean ± SEM. **(D)** Representative images from Sca1^**+**^/Tcf21^lin^ trilineage differentiation. Adipogenic differentiation was analyzed at 10 days by Perilipin staining, chondrogenic differentiation at 14 days by Sox9 staining, and osteogenic differentiation at 21 days by Osterix staining. **(E)** Immunofluorescence staining of SMA or SF1 in the Tcf21^lin^ clonal cells differentiated into Leydig or myoid cells. **(F)** Serum testosterone levels measured from Sca1^+^/cKit^-^ and cKit^+^/Sca1^-^ cells at 3, 9, and 15 days of culture in the presence of LH. Data are presented as mean ±SEM. **(G)** Serum testosterone levels measured from Sca1^+^/cKit^-^ and cKit^+^/Sca1^-^ cells at 4, 7, and 14 days of culture in the absence of LH. Data are presented as mean ± SEM. **(H)** Rank correlation of the seven centroids from the *in vitro* Leydig time-course differentiation with 6 cluster centroids derived from previously published adult mouse somatic cells (Green et al., 2018) and **(I)** previously published Nr5a1^+^GFP^+^ progenitor cells from E10.5 – E16.5 male mouse gonads (Stevant et al., 2018).

**Supplemental Figure 3: The fetal Tcf21^lin^ population is a bipotential somatic progenitor. (A)** *Tcf21^+/+^:R26R^tdTom^* or *Tcf21^mCrem^:R26R^tdTom^* time pregnant females (E10.5) were treated with Tamoxifen (left panels) or corn oil (right panels), respectively, confirming specificity and tightness of Cre expression. (**B)** Co-immunostaining of the Tcf21^lin^ with germ cell marker Vasa **(Ddx4; green)** in the fetal testis (collected at different injection timepoints). **(C-H)** Quantification of the percentage of various somatic cells co-labeled with Tcf21^lin^ (black bars) and percentage of cells in Tcf21^lin^-positive population co-expressing somatic markers (gray bars) at different embryonic days. **(I, J)** Colocalization of Tcf21^lin^ and Gli1-egfp **(I)**, Pdgfra^eGFP^ **(J)**, and the nuclear counterstain DAPI. **(K)** Tcf21^lin^ cells in the E11.5 fetal ovary are present in the coelomic epithelium and mesonephros. Overlaying the Tcf21^lin^ (Tdtom) cells with WT1+ cells **(Green)** for **(L)** E10.5-E12.5 Tcf21^lin^ cells label somatic cells broadly in the fetal ovary at E17.5. **(M-P**) Overlay of the Tcf21^lin^ with granulosa cell marker Foxl2 **(green; M)**, the interstitial cell marker CoupTFII (**Nr2F2, green; N**); the smooth muscle marker SMA **(green; O)**; the germ cell marker Vasa **(Ddx4, green; P)**. **(Q)** The E10.5 fetal Tcf21^lin^ contributes to multiple somatic cell types in the adult ovary. In all panels the nuclear counterstain is DAPI. Scale bars: 20 μm.

**Supplemental Figure 4: Peritubular myoid cells can regenerate after Diphtheria toxin treatment.**

**(A)** Schematic representation of the genetic method used to ablate myoid cells. **(B-E)** Efficacy of diphtheria toxin treatment on MYH11^cre-egfp^; Rosa26^iDTR/+^ or MYH11^cre-egfp^; Rosa26^+/+^ animals. **(B)** Representative images of apoptotic myoid cell death by TUNEL staining in testes collected 12hpfi or 4dpfi of DTX, **(C)** Representative H&E staining at 12hpfi or 4dpfi of DTX. **(D)** BrdU^+^ cells are detected in peritubular cells of MYH11^cre-egfp^:Rosa26^iDTR^ testes. Representative images of BrdU^+^/SMA^+^ (yellow arrows) or BrdU^+^/SMA^-^ (white arrows)_at 4dpfi in MYH11^cre-egfp^:Rosa26^+/+^ or Myh11^cre-egfp^:Rosa26^iDTR/+^ animals. **(E)** Quantification of BrdU/SMA coexpression at 12hpfi or 4dpfi of DTX. Scale bars: 20 μm.

**Supplemental Figure 5: Tcf21^lin^ cells home to the seminiferous tubule basement membrane after injury. (A)** Schematic representation of the Tcf21^lin^ transplant following genetic ablation of myoid cells in MYH11^cre-egfp^:Rosa26^iDTR/+^ animals. (**B**) Representative immunofluorescence images of Tcf21^lin^ in MYH11^cre-egfp^:Rosa26^iDTR/+^ animals at 24hpt. Scale bars: 20 μm.

**Supplemental Figure 6: The juvenile Tcf21^lin^ cells contribute to testis homeostasis.**

**(A)** 5-week old *Tcf21^+/+^:R26R^tdTom^* or *Tcf21^mCrem^:R26R^tdTom^* males were injected with 3 doses of tamoxifen. The contribution of the Tcf21^Lin^ was analyzed one week or one year following Tamoxifen treatment. Images illustrate the colocalization of Tcf21^lin^ with the Leydig cell marker SF1 (green). DAPI is used as the nuclear counterstain (white). Dotted lines indicate seminiferous tubules. Scale bars: 20 μm.

## Supplemental Table Legends

**Table S1. Markers for Somatic Cell Types (Related to Figure 1)**

A. Markers for 6 Somatic Cell Types (Related to Figure 1A and 1B)

B. Markers for 4 Somatic Cell Types (Related to Figure 1C and 1D)

C. GO Term Enrichment for TCF21+ Int Population (Related to Figure 1D)

**Table S2. Markers for In-vitro Leydig Differentiation (Related to Figure 2)**

A. Markers for 7 clusters of in-vitro Leydig differentiation

B. Gene expression centroids of 7 clusters of in-vitro Leydig differentiation

**Table S3. Markers for Somatic Cell Types (Related to Figure 6)**

A. Interaction scores between Tcf21+ Interstitial cells and all other somatic cells in adult mouse testis, calculated by multiplying the centroid values for each cell type from scRNAseq dataset from Green et al. Top 5% of scores are shown, others are truncated to 0.

B. Interaction scores between Tcf21+ Interstitial cells and spermatogonia, calculated in the same way as A.

