## SupplementalFigures 1-7 for "Tcf21^+^ mesenchymal cells contribute to testis somatic cell development, homeostasis, and regeneration"

#### Supplemental Figure 1

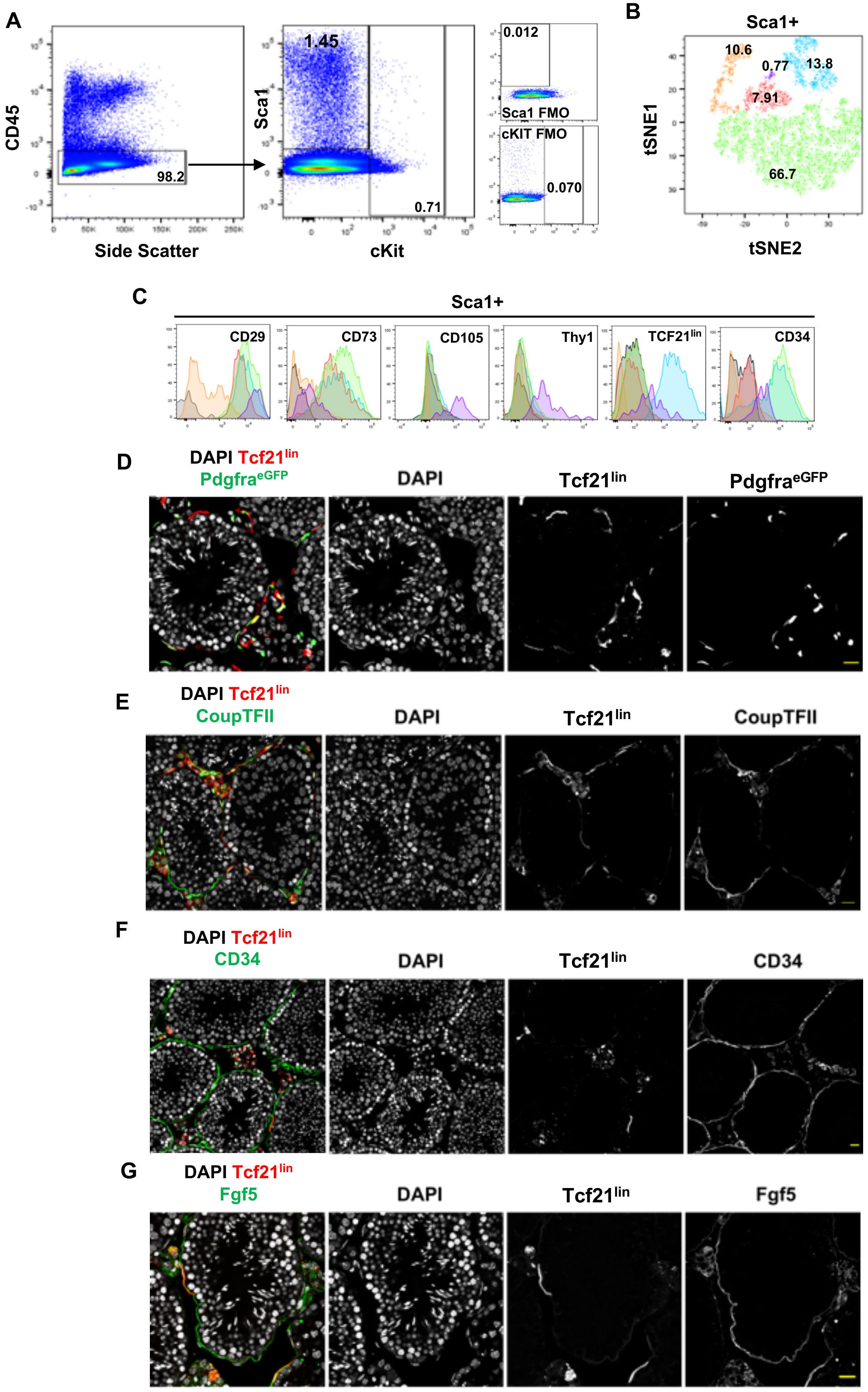

#### Supplemental Figure 2

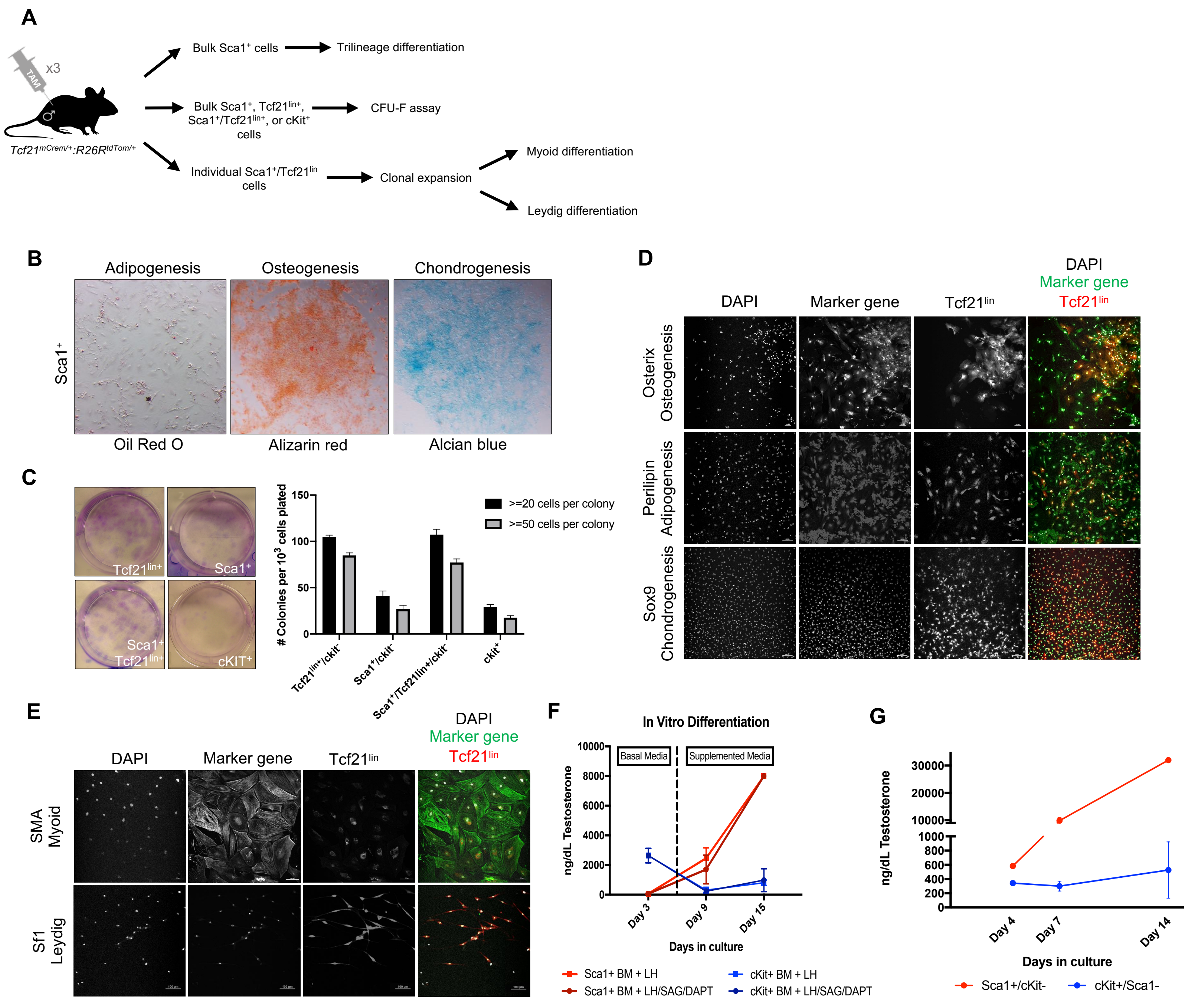


**H**

| | InnateLymphoid | Macrophage | Endothelial | Myoid | *Tcf21*+Int | Leydig | Sertoli |
|---|---|---|---|---|---|---|---|
| 1.Endothelial | 0.26 | 0.36 | 0.89 | 0.59 | 0.54 | 0.43 | 0.35 |
| 2.IntProg | 0.32 | 0.61 | 0.58 | 0.63 | 0.98 | 0.73 | 0.41 |
| 3.ProlifProg | 0.39 | 0.50 | 0.56 | 0.63 | 0.74 | 0.58 | 0.37 |
| 4.ECMmyofibroblast | 0.33 | 0.50 | 0.56 | 0.66 | 0.77 | 0.62 | 0.41 |
| 5.DiffMyofibroblast | 0.37 | 0.53 | 0.58 | 0.67 | 0.80 | 0.61 | 0.36 |
| 6.DiffLeydig | 0.28 | 0.54 | 0.51 | 0.61 | 0.85 | 0.73 | 0.42 |
| 7.Leydig | 0.24 | 0.47 | 0.44 | 0.53 | 0.76 | 0.84 | 0.47 |

**I**

| | C1.Endothelial | C2.EarlyProg | C3.IntProg | C4.Pre.Sertoli | C5.FetalLeydig | C6.Sertoli |
|---|---|---|---|---|---|---|
| 1.Endothelial | 0.49 | 0.35 | 0.39 | 0.35 | 0.35 | 0.36 |
| 2.IntProg | 0.42 | 0.41 | 0.56 | 0.40 | 0.46 | 0.39 |
| 3.ProlifProg | 0.46 | 0.54 | 0.64 | 0.39 | 0.48 | 0.45 |
| 4.ECMmyofibroblast | 0.46 | 0.49 | 0.58 | 0.43 | 0.45 | 0.41 |
| 5.DiffMyofibroblast | 0.46 | 0.49 | 0.60 | 0.41 | 0.45 | 0.40 |
| 6.DiffLeydig | 0.43 | 0.46 | 0.61 | 0.45 | 0.53 | 0.46 |
| 7.Leydig | 0.39 | 0.42 | 0.55 | 0.49 | 0.58 | 0.50 |

### Supplemental Figure 3

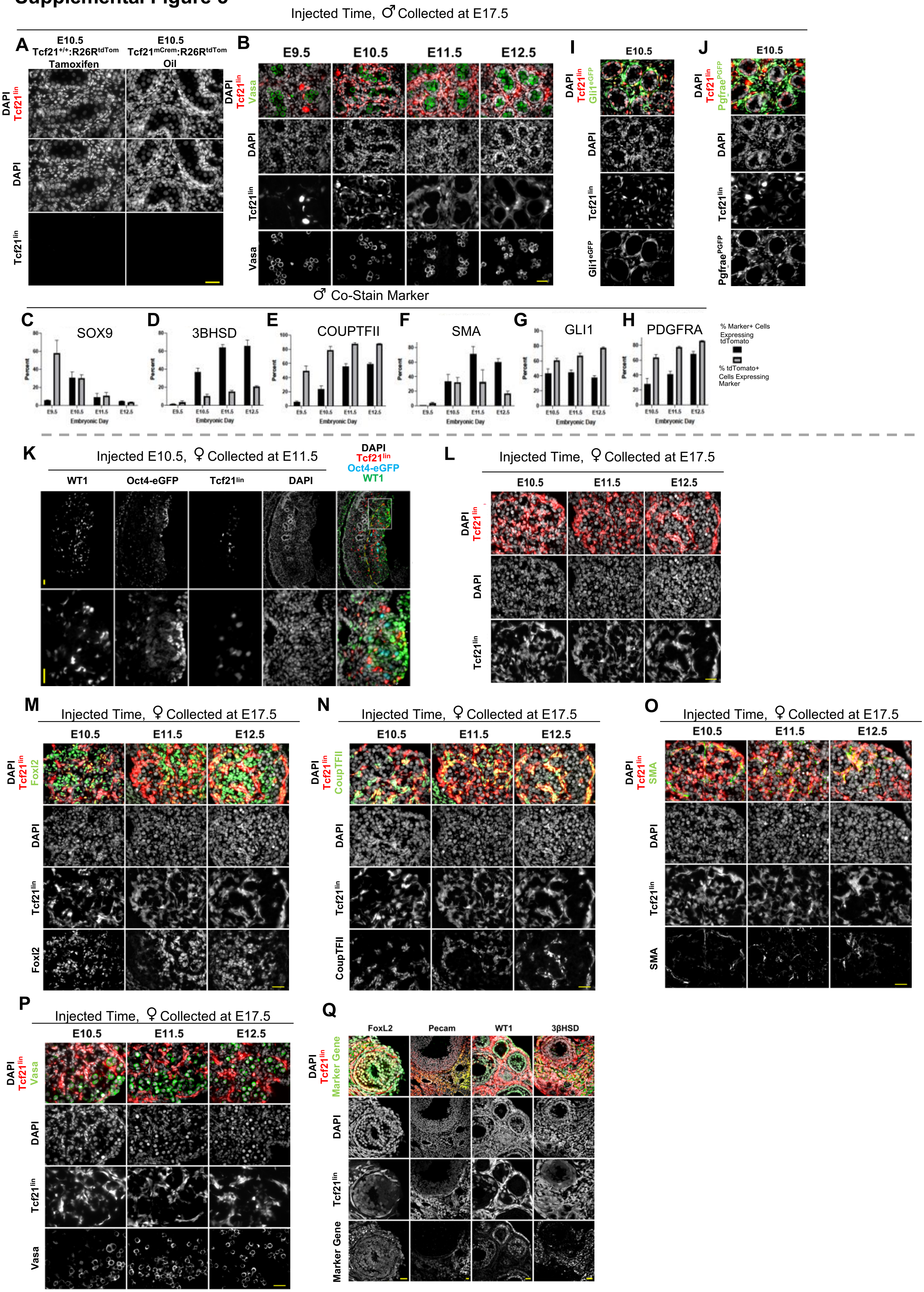

### Supplemental Figure 4

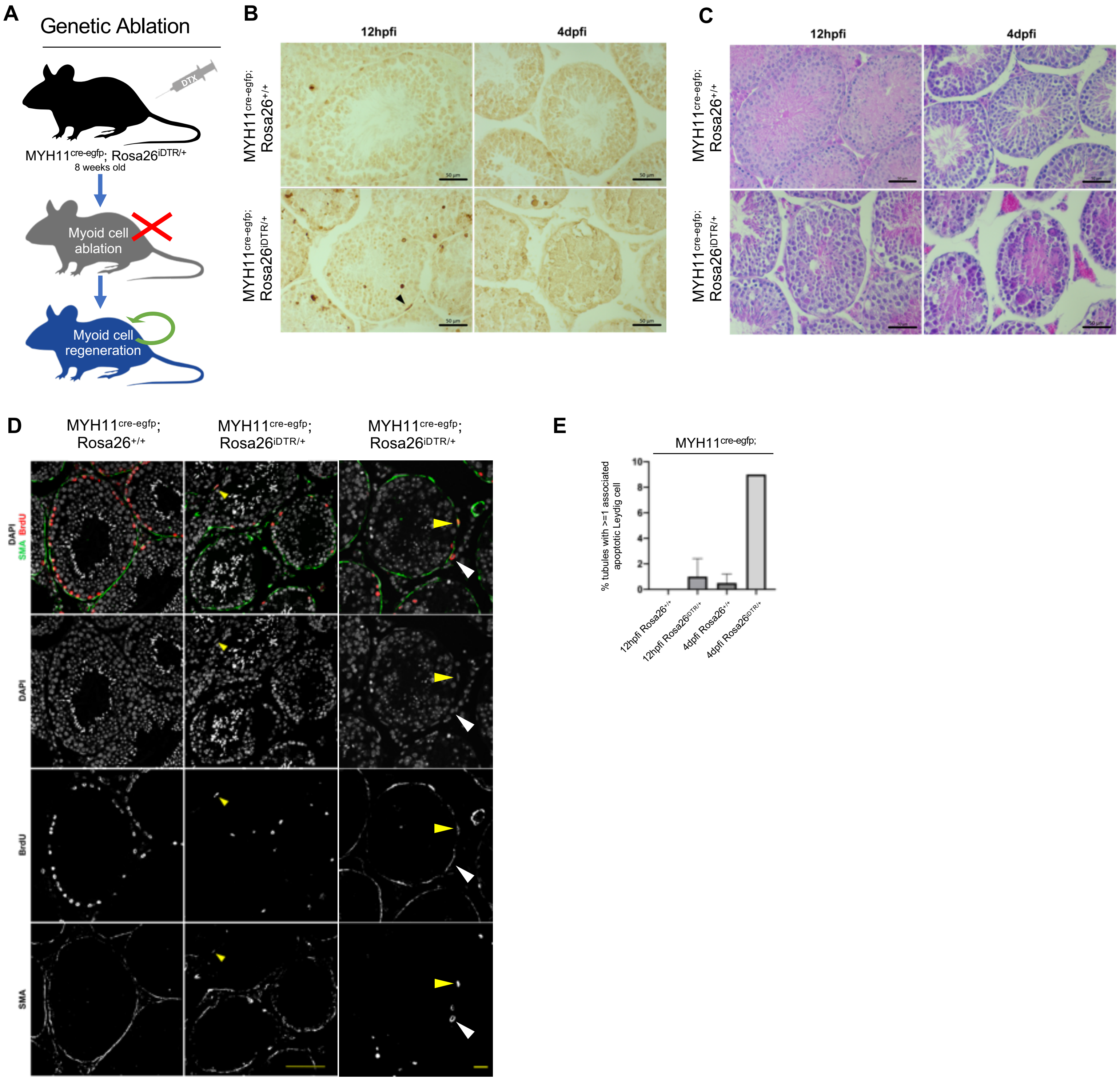

### Supplemental Figure 5

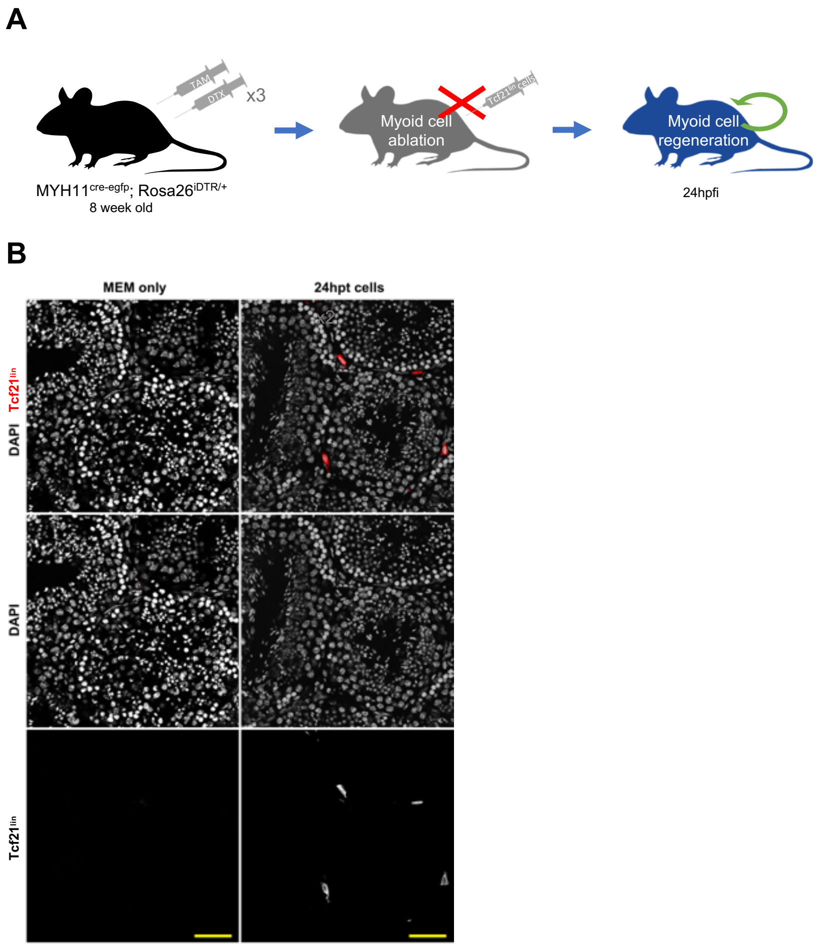

**Supplemental Figure 6**

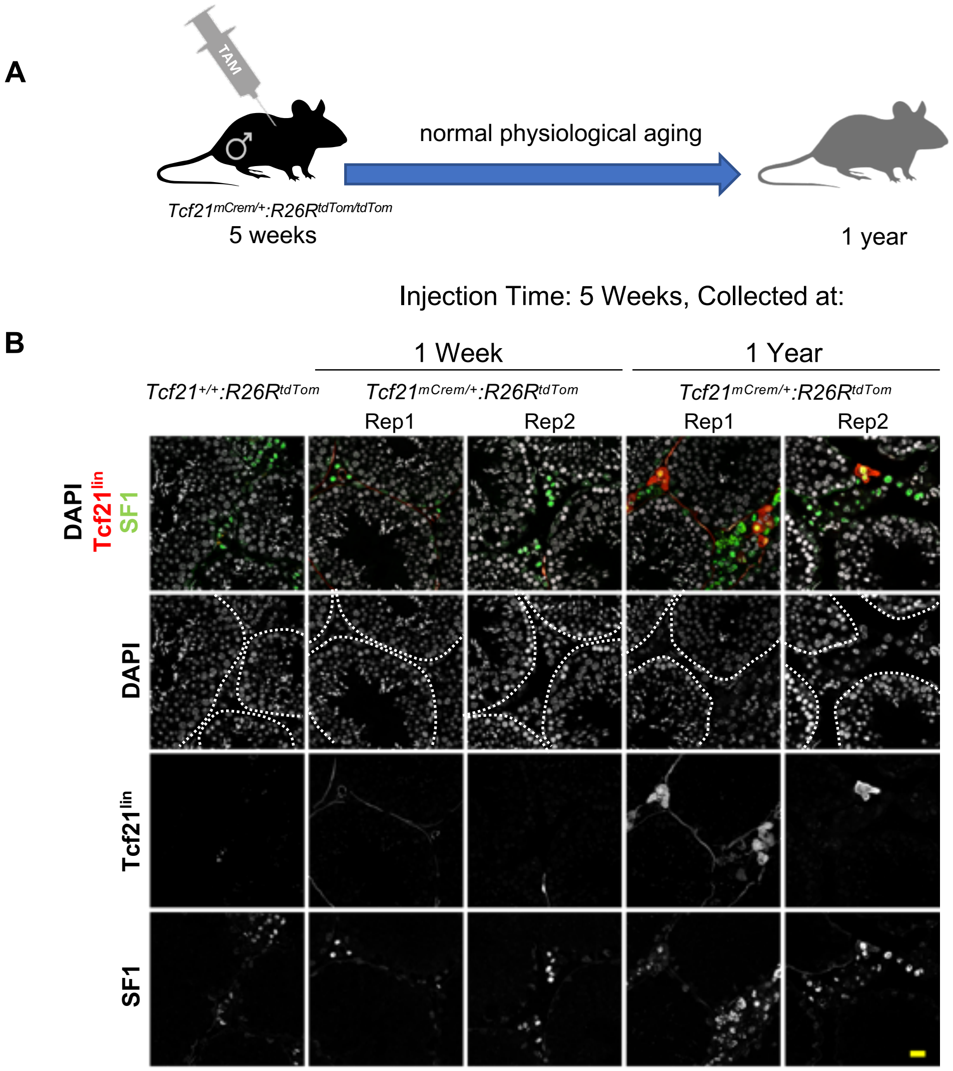
